## Supplemental Information for "Reconstitution of Early Paclitaxel Biosynthetic Network"

#### Table of Contents

**Supplementary Tables** 2
**Supplementary Figures** 24
**Supplementary References** 68

### Supplementary Tables

**Supplementary Table 1. Reported product profile of T5αH in different systems.**

| Host | iso-OCT | OCT | Taxadien-5α-ol | Diols | Others | Year/Reference |
|---|---|---|---|---|---|---|
| *S. cerevisiae* | | | ✓ | ✓ | | 2004[1] (Original discovery of T5αH) |
| *Spodoptera fugiperd* (armyworm) | | | ✓ | | | |
| *N. sylvestris* trichome | | ✓ | | | | 2008[2] |
| *S. cerevisiae* microsome | | ✓ | | | | |
| *E. coli* | ✓ | ✓ | ? | | | 2010[3] |
| *E. coli* | ✓ | ✓ | ✓ | ✓ | ✓ | 2014[4] |
| *E. coli* (TS) + *S. cerevisiae* (T5αH) consortium | | ✓ | ✓ | | ✓ | 2015[5] |
| *In vitro* lipid nanodisc | | ✓ | ✓ | ✓ | ✓ | 2016[6] |
| Yarrowia lipolytica | ✓ | ✓ | ✓ | ✓ | ✓ | 2016[7] |
| *E. coli* | ✓ | ✓ | ✓ | | ✓ | 2018[8] |
| *N. benthamiana* | ✓ | ✓ | ✓ | | | 2019[9] |
| *S. cerevisiae* | ✓ | ✓ | ✓ | ✓ | ✓ | 2021[10] |
| *S. cerevisiae* microsome | ✓ | ✓ | ✓ | | | 2021[11] |
| *S. cerevisiae* | ✓ | ✓ | ✓ | ✓ | | 2022[12] |

**Supplementary Table 2. Mass spectra of all compounds in this study.**

| Compound | Molecular formula | Calc. mass | Retention time (min) | Mass spectra |
|---|---|---|---|---|
| taxadiene (**1**) | $C_{20}H_{32}$ | 272.2504 | 10.61 (GCMS) | GCMS: 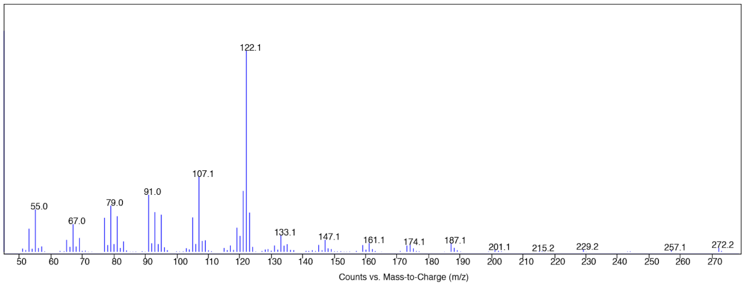  |
| taxadien-5α-ol (**2**) | $C_{20}H_{32}O$ | 288.2453 | 12.17 (GCMS) | GCMS: 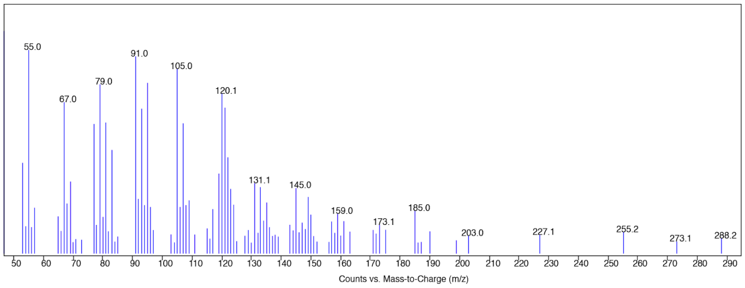  |
| OCT (**3**) | $C_{20}H_{32}O$ | 288.2453 | 11.89 (GCMS) | GCMS: 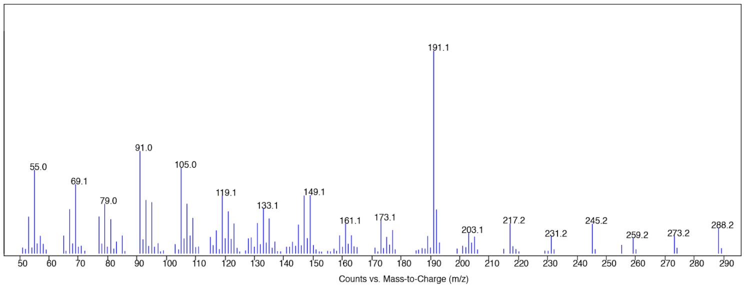  |
| iso-OCT (**4**) | $C_{20}H_{32}O$ | 288.2453 | 11.33 (GCMS) | GCMS: 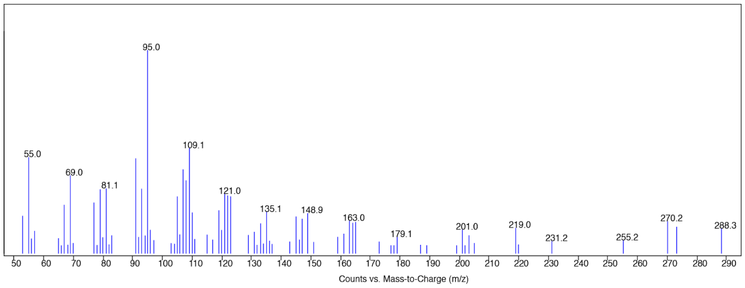  |
| **5** | $C_{20}H_{32}O$ | 288.2453 | 12.90 (GCMS) | GCMS: |

<table>
<tr><td></td><td></td><td></td><td></td><td>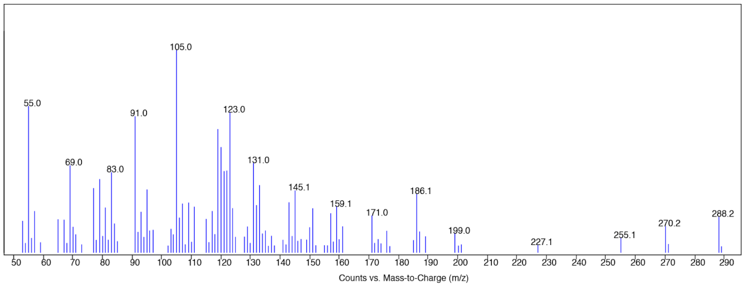
</td></tr>
<tr><td>6</td><td>$C_{20}H_{32}O_2$</td><td>304.2402</td><td>13.90 (GCMS)</td><td>GCMS:<br>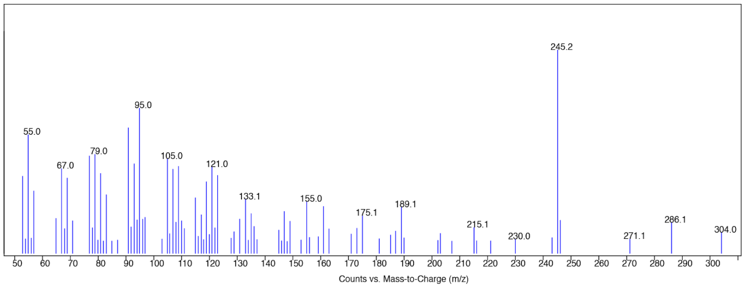
</td></tr>
<tr><td>acetylated 6</td><td>$C_{22}H_{34}O_3$</td><td>346.2508</td><td>15.02 (GCMS)</td><td>GCMS:<br>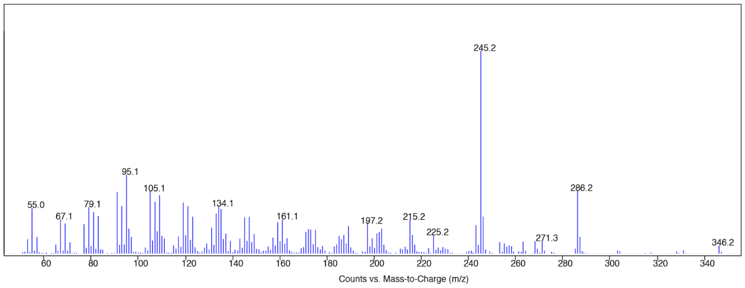
</td></tr>
<tr><td>7</td><td>$C_{20}H_{32}O_2$</td><td>304.2402</td><td>14.49 (GCMS)</td><td>GCMS:<br>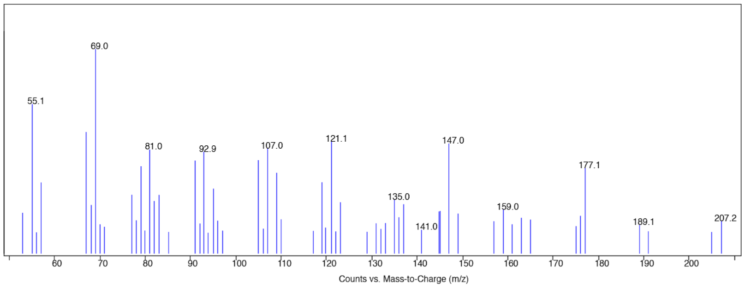
</td></tr>
<tr><td>acetylated 7</td><td>$C_{22}H_{34}O_3$</td><td>346.2508</td><td>14.83 (GCMS)</td><td>GCMS:</td></tr>
</table>

| | | | | |
|---|---|---|---|---|
| | | | | 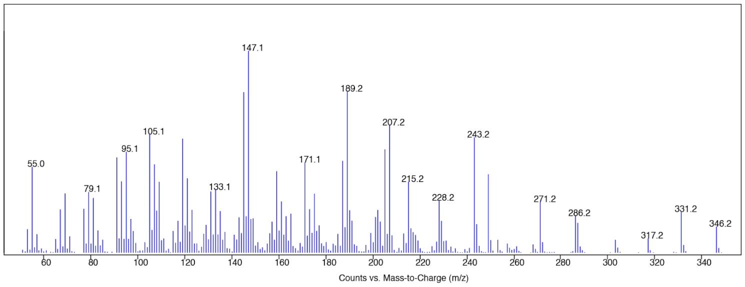 |
| **8** | $C_{20}H_{34}O_2$ | 306.2559 | 15.80 (GCMS) | GCMS: 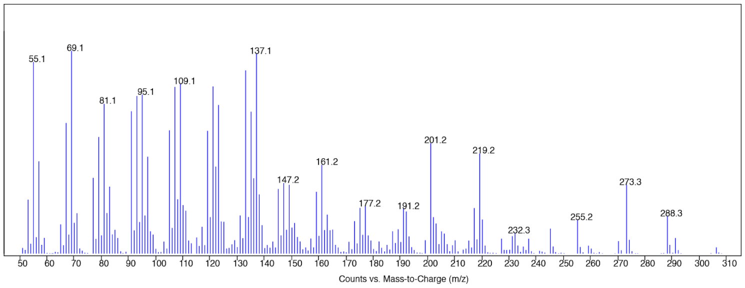 |
| **9** | $C_{20}H_{32}O_2$ | 304.2402 | 13.82 (GCMS) | GCMS: 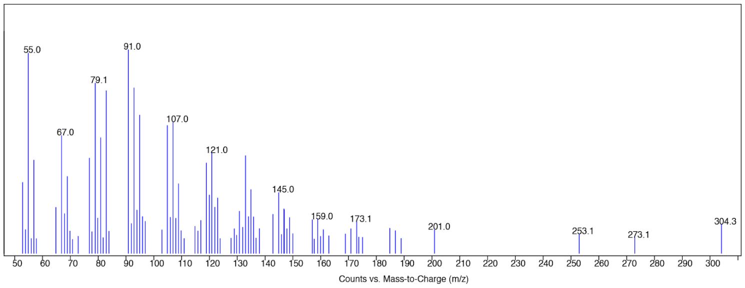 |
| **10** (acetylated **9**) | $C_{22}H_{34}O_3$ | 346.2508 | 14.80 (GCMS) | GCMS: 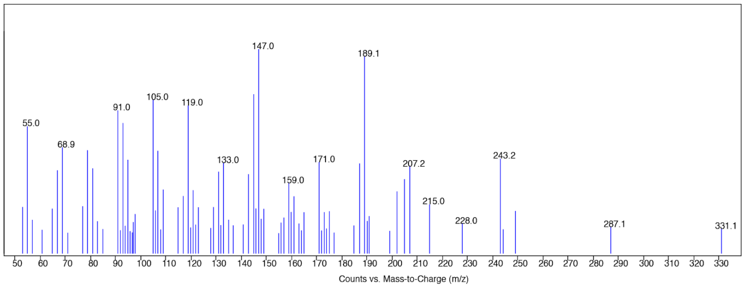 |
| taxadien-5α-acetoxy (**11**) | $C_{22}H_{34}O_2$ | 330.2559 | 13.77 (GCMS) | GCMS: |

| | | | | |
|---|---|---|---|---|
| | | | | 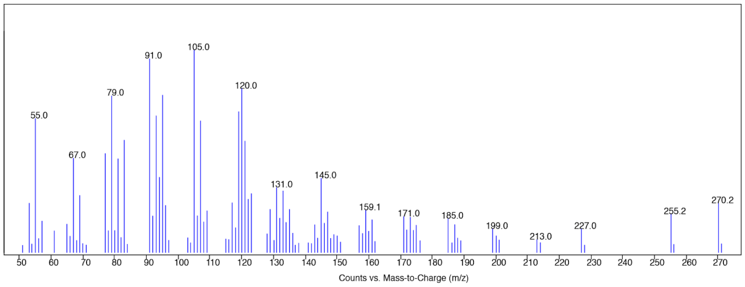  |
| taxadien-5α-acetoxy-10β-ol (**12**) | $C_{22}H_{34}O_3$ | 346.2508 | 15.79 (GCMS) | GCMS: 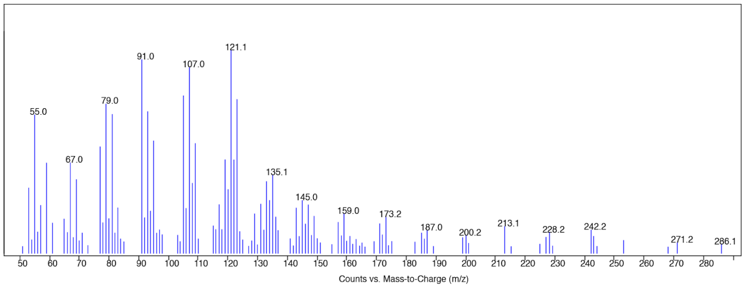  |
| taxadien-5α, 10β-diacetox y (**13**) | $C_{24}H_{36}O_4$ | 388.2614 | 16.06 (GCMS) | GCMS: 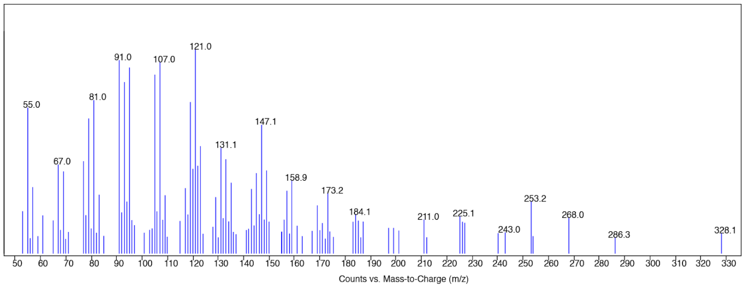  |
| taxadien-5α, 10β-diacetox y-13α-ol (**14**) | $C_{24}H_{36}O_5$ | 404.2563 | 17.91 (GCMS), 8.66 (LCMS) | GCMS: 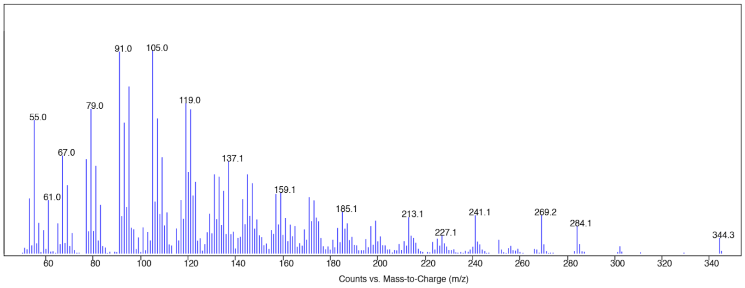  LCMS: 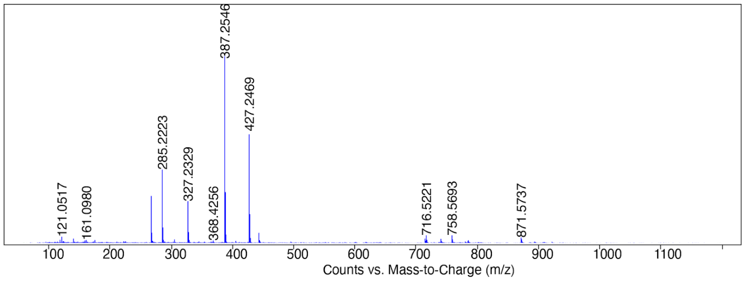  |

| | | | | |
|---|---|---|---|---|
| taxadien-5α, 10β-diacetoxy-13-one (**15**) | $C_{24}H_{34}O_5$ | 402.2406 | 17.50 (GCMS), 8.32 (LCMS) | GCMS: 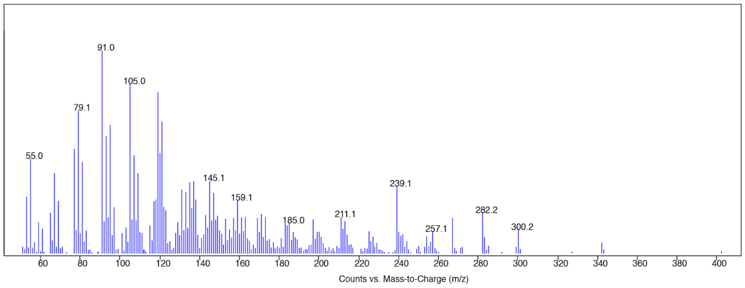  LCMS: 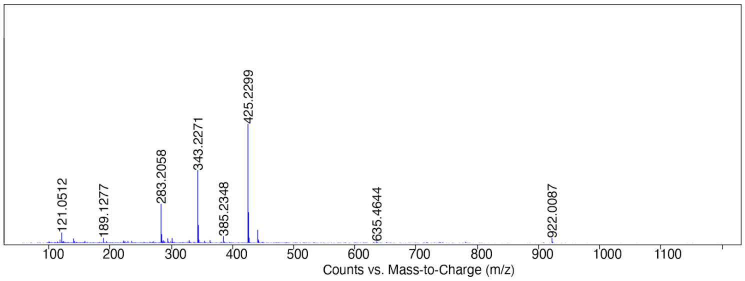  |
| taxadien-5α, 13α-diol (**16**) | $C_{20}H_{32}O_2$ | 304.2402 | 10.17 (LCMS) | LCMS: 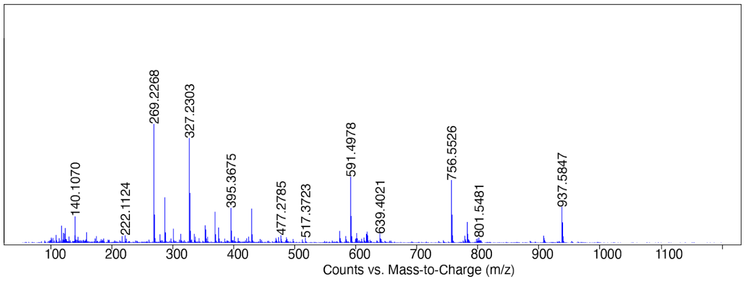  |
| Taxadien-5α-ol-13-one (**17**) | $C_{20}H_{30}O_2$ | 302.2246 | 9.40 (LCMS) | LCMS: 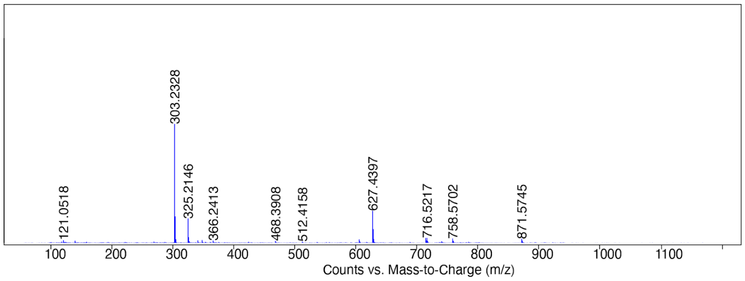  |
| taxadien-5α-acetoxy-13α-ol (**18**) | $C_{22}H_{34}O_3$ | 346.2508 | 15.83 (GCMS) | GCMS: 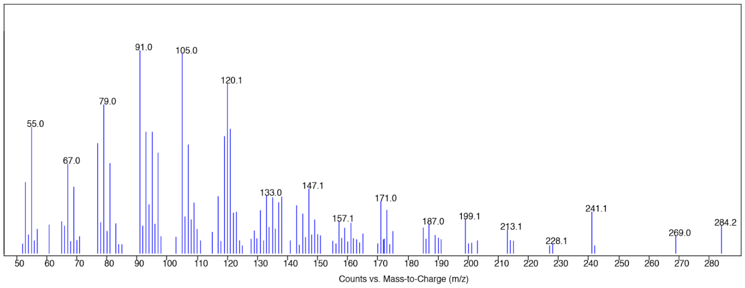  |

| | | | | |
|---|---|---|---|---|
| taxadien-5α-acetoxy-13-one (**19**) | $C_{22}H_{32}O_3$ | 344.2351 | 9.92 (LCMS) | LCMS: 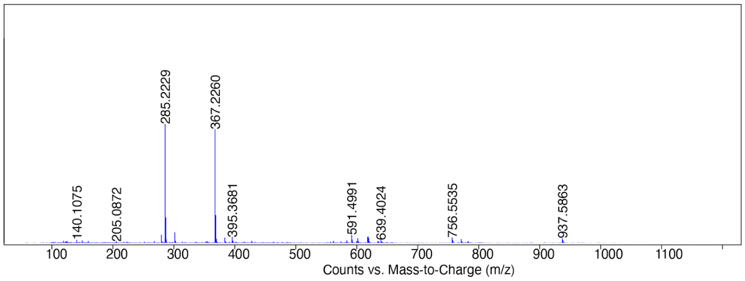  |
| taxadien-5α-acetoxy-10β, 13α-diol (**20**) | $C_{22}H_{34}O_4$ | 362.2457 | 17.66 (GCMS) | GCMS: 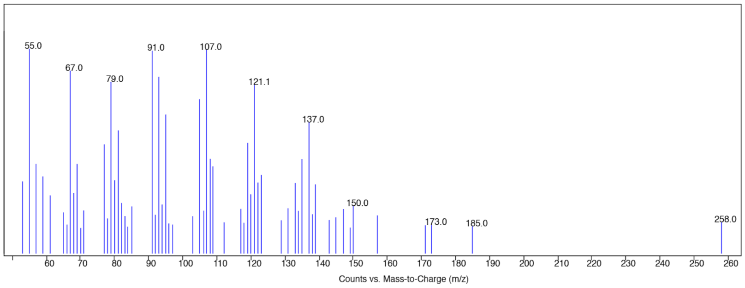  |
| taxadien-5α-acetoxy-10β-ol-13-one (**21**) | $C_{22}H_{32}O_4$ | 360.2301 | 6.69 (LCMS) | LCMS: 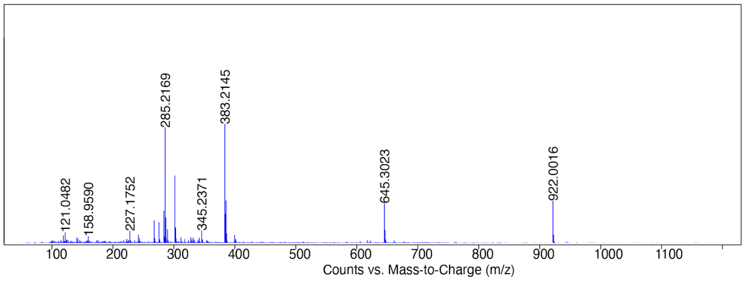  |

**Supplementary Table 3. Summary of compound purification in this study.**

| Name | Source | Scale | Yield (mg) | Column condition | NMR |
|---|---|---|---|---|---|
| **1** | Yeast expressing TS | 400 mL culture | 15.5 | 20 g silica column (isocratic 100% hexane) | Supplementary Figure 1~3 |
| **2** | synthesis | 15.5 mg of **1** | 1.8 | 200 mg silica column (hexane:diethyl ether = 5:1) | Supplementary Figure 4-5 |
| **3** | *N. benthamiana* transiently expressing tHMGR+GGPPS+TS1/2+T5αH+TAX19 | 58 * 6-weeks old plants (73.47 g DW) | 0.4 | 1. 250 g silica column (EA:Hex = 3:7, 2 L; EA:Hex = 4:6, 1 L; EA:Hex = 1:1, 1L; EA:Hex = 6:4, 1L; EA:Hex = 8:2, 1L) 2. Biotage C18 6 g column (50% 5 CV, 50-65% over 45 CV) | Supplementary Figure 6-7 |
| **5** | *N. benthamiana* transiently expressing tHMGR+GGPPS+TS1/2+(NOS)T5αH+TAT+(NOS)T10βH+DBAT+(NOS)T13αH | 26 * 4-weeks old plants (14.31 g DW) | 0.4 | 1. 100 g silica column (EA:Hex = 2:8, 1 L; EA:Hex = 3:7, 1 L; EA:Hex = 1:1, 1 L) 2. Biotage C18 6 g column (50% 3 CV, 50-70% over 20 CV) | Supplementary Figure 8~13 Supplementary Table 3 |
| acetylated **6** | *N. benthamiana* transiently expressing tHMGR+GGPPS+TS1/2+T5αH+TAX19 | 58 * 6-weeks old plants (73.47 g DW) | 1.5 | 1. 250 g silica column (EA:Hex = 3:7, 2 L; EA:Hex = 4:6, 1 L; EA:Hex = 1:1, 1L; EA:Hex = 6:4, 1L; EA:Hex = 8:2, 1L) 2. Biotage C18 6 g column (50% 5 CV, 50-65% over 45 CV) | Supplementary Figure 14~20 Supplementary Table 4 |
| acetylated **7** | *N. benthamiana* transiently expressing tHMGR+GGPPS+TS1/2+T5αH+TAX19 | 58 * 6-weeks old plants (73.47 g DW) | 1.7 | 1. 250 g silica column (EA:Hex = 3:7, 2 L; EA:Hex = 4:6, 1 L; EA:Hex = 1:1, 1L; EA:Hex = 6:4, 1L; EA:Hex = 8:2, 1L) 2. Biotage C18 6 g column (50% 5 CV, 50-65% over 45 CV) | Supplementary Figure 21~25 Supplementary Table 5 |
| **8** | *S.cerevisiae* expressing TS+T5αH | 4 L culture | 1.8 | 1. 25 g Biotage silica column (100% Hex 10 CV, 100-85% Hex/EA over 15 CV) 2. Biotage C18 12 g column (40% 5 CV, 40-50% over 30 CV) | Supplementary Figure 26~32 Supplementary Table 6 |

| | | | | | |
|---|---|---|---|---|---|
| **14** | *N. benthamiana* transiently expressing tHMGR+GGPPS+TS1/2+(NOS)T5αH+TAT+(NOS)T10βH+DBAT+(NOS)T13αH | 26 * 4-weeks old plants (14.31 g DW) | 0.9 | 1. 100 g silica column (EA:Hex = 2:8, 1 L; EA:Hex = 3:7, 1 L; EA:Hex = 1:1, 1 L)<br>2. Biotage C18 6 g column (50% 3 CV, 50-70% over 20 CV) | Supplementary Figure 33~38 Supplementary Table 8 |
| **15** | *N. benthamiana* transiently expressing tHMGR+GGPPS+TS1/2+(NOS)T5αH+TAT+(NOS)T10βH+DBAT+(NOS)T13αH | 26 * 4-weeks old plants (14.31 g DW) | 0.6 | 1. 100 g silica column (EA:Hex = 2:8, 1 L, EA:Hex = 3:7, 1 L, EA:Hex = 1:1, 1 L)<br>2. Biotage C18 6 g column (50% 3 CV, 50-65% over 15 CV) | Supplementary Figure 39~44 Supplementary Table 8 |

Solvent system for biotage Sfär C18 D Duo 100 Å 30 μm column: A = acetonitrile, B = water. Percentage of solvent B is listed. TAX19 is a TAT homolog that has 5α-*O*-acetylation activity on taxadien-5α-ol but shows different regioselectivity to TAT on taxusin-tetraol substrate compared to TAT.[14]
DW: dry weight, EA: ethyl acetate, Hex: hexane, CV: column volume

**Supplementary Table 4. $^{13}C$ & $^{1}H$ δ assignments of mono-oxidized taxadiene 5.**

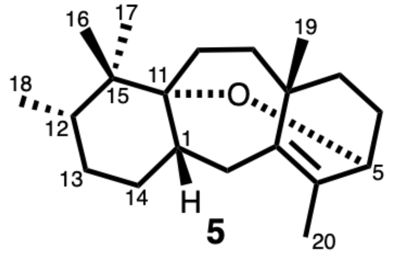


| Carbon | δ $^{13}C$ (ppm) | δ $^{1}H$ (mult.; *J* in Hz) |
|---|---|---|
| **1** | 42.86 | 1.97 (dt; 6.0, 5.5) |
| **2** | 27.19 | 2.23 (dd; 6.0, 13.6),<br>2.42 (d; 13.6) |
| **3** | 140.7 | - |
| **4** | 131.7 | - |
| **5** | 70.2 | 3.80 (d; 4.3) |
| **6** | 29.21 | 1.77 (ddt; 3.5, 4.3, 14.4),<br>2.08 (tt; 4.3, 14.4) |
| **7** | 31.95 | 1.10 (dt; 14.6, 3.5),<br>1.86 (dt; 13.6, 3.5) |
| **8** | 38.70 | - |
| **9** | 35.41 | 1.12 (dt; 12.9, 3.5), 2.18 (m) |
| **10** | 24.51 | 1.49 (m) |
| **11** | 77.14 | - |
| **12** | 37.56 | 1.43 (m) |
| **13** | 30.34 | 1.27 (m), 1.33 (m) |
| **14** | 27.73 | 1.40 (m), 1.54 (m) |
| **15** | 42.88 | - |
| **16** | 16.16 | 0.86 (s) |
| **17** | 23.36 | 0.84 (s) |
| **18** | 16.32 | 0.78 (d; 6.7) |
| **19** | 28.22 | 0.93 (s) |

| | | |
|---|---|---|
| **20** | 19.95 | 1.74 (s) |

s = singlet, d = doublet, dd = doublet of doublets, dt = doublet of triplets, t = triplet, q = quartet, quint = quintet, m = multiplet

**Supplementary Table 5. $^{13}C$ & $^{1}H$ δ assignments of acetylated 6**

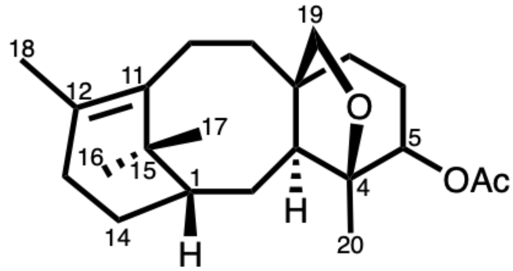


acetylated **6**

| Carbon | δ $^{13}C$ (ppm) | δ $^{1}H$ (mult.; *J* in Hz) |
|---|---|---|
| **1** | 43.05 | 1.78 (m) |
| **2** | 27.29 | 1.49 (m) |
| **3** | 42.16 | 2.25 (dd; 2.2, 7.6) |
| **4** | 86.1 | - |
| **5** | 77.75b | 4.68 (m) |
| **6** | 25.18 | 1.63 (m), 1.99a (m) |
| **7** | 32.54 | 1.19, 1.99b (m) |
| **8** | 47.26 | - |
| **9** | 33.42 | 1.27 (m), 2.00 (m) |
| **10** | 24.03 | 2.09 (d; 3.5),<br>2.61 (dt; 5.3, 13.4) |
| **11** | 136.40 | - |
| **12** | 129.83 | - |
| **13** | 29.97 | 1.96 (m), 2.37 (m) |
| **14** | 23.02 | 1.36 (m), 2.15 (m) |
| **15** | 38.63 | - |
| **16** | 30.83 | 1.05 (s) |
| **17** | 25.26 | 1.30 (s) |
| **18** | 21.75 | 1.76 (s) |
| **19** | 77.75a | 3.56 (dd; 1.7, 8.0),<br>3.61 (d; 8.0) |

| | | |
|---|---|---|
| **20** | 21.43 | 1.13 (s) |
| **$-\underline{C}O_2Me$** | 169.80 | - |
| **$-CO_2\underline{Me}$** | 21.36 | 2.07 |

**Supplementary Table 6. $^{13}C$ & $^{1}H$ δ assignments of acetylated 7**

| | 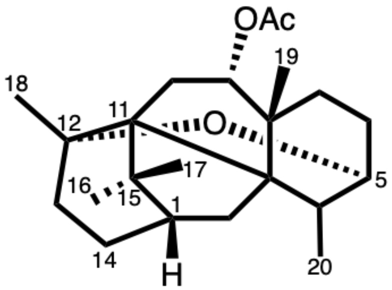 acetylated **7** | | 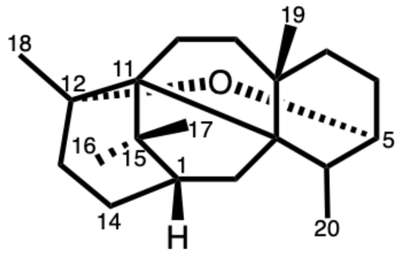 OCT (**3**) | |
|---|---|---|---|---|
| **Carbon** | **δ $^{13}C$ (ppm)** | **δ $^{1}H$ (mult.; *J* in Hz)** | **δ $^{13}C$ (ppm)**[2] | **δ $^{1}H$ (mult.; *J* in Hz)**[2] |
| **1** | 44.58 | 1.74 (m) | 45.9 | 1.71 (dd; 5.0, 8.4) |
| **2** | N.A. | 1.41 (m),<br>2.24 (dd; 5.2, 13.0) | 39.1 | 1.33 (d; 12.9),<br>2.21 (dd; 5.0, 12.9) |
| **3** | 51.12 | - | 53.3 | - |
| **4** | N.A. | 2.51 (qd; 7.0, 3.5) | 37.1 | 2.47 (qd; 7.0, 3.4) |
| **5** | 69.45 | 4.01 (dd; 3.6, 8.7) | 69.8 | 3.97 (dd; 3.4, 9.1) |
| **6** | N.A. | 2.06 (m) | 30.2 | 1.83 (m), 2.04 (m) |
| **7** | N.A. | N.A. | 37.5 | 1.36 (d; 10.9), 1.83 (m) |
| **8** | 43.78 | - | 42.7 | - |
| **9** | 82.18 | 4.98 (dd; 7.5, 11.5) | 47.3 | 1.53 (dd; 8.8, 12.7), 1.82 (m) |
| **10** | N.A. | 1.82 (dd; 7.9, 12.4),<br>1.93 (t, 12.0) | 30.2 | 1.31 (dd; 3.6, 9.6),<br>1.38 (ddd; 1.0, 3.6, 10.9) |
| **11** | 60.00 | - | 66.0 | - |
| **12** | 79.45 | - | 80.5 | - |
| **13** | N.A. | N.A. | 36.4 | 1.84 (m), 1.98 (m) |
| **14** | N.A. | N.A. | 28.1 | 1.62 (ddd; 5.2, 11.2, 14.3),<br>2.01 (m) |
| **15** | N.A. | - | 46.0 | - |
| **16** | 29.22 | 1.03 (s) | 28.6 | 0.93 (brs) |
| **17** | 26.83 | 1.07 (s) | 26.9 | 1.01 (brs) |

| | | | | |
|---|---|---|---|---|
| **18** | 30.15 | 1.19 (s) | 30.3 | 1.19 (s) |
| **19** | 26.34 | 1.02 (s) | 28.0 | 1.04 (s) |
| **20** | 15.28 | 1.19 (d; 7.0) | 15.2 | 1.13 (d; 7.0) |
| **$-\underline{C}O_2Me$** | 170.81 | - | - | - |
| **$-CO_2\underline{Me}$** | 21.46 | 2.07 | - | - |

Chemical shifts of OCT from reference[2] are shown as comparison.
* N.A.:  chemical shift not assigned due to insufficient information

**Supplementary Table 7. $^{13}$C & $^{1}$H δ assignments of 8.**

| | 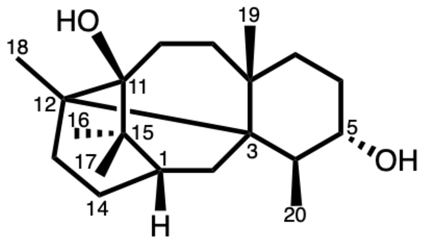 **8** | |
|---|---|---|
| **Carbon** | **δ $^{13}$C (ppm)** | **δ $^{1}$H (mult.; *J* in Hz)** |
| **1** | 39.51 | 1.22b (m) |
| **2** | 30.09 | 1.63 (m), 1.82 (dd; 4.0, 15.1) |
| **3** | 45.77 | - |
| **4** | 40.66 | 1.74 (m) |
| **5** | 73.94 | 3.87 (ddd; 6.8, 9.1, 12.5) |
| **6** | 28.65 | 1.65 (m), 1.96 (dddd; 2.0, 6.8, 11.7, 13.8) |
| **7** | 32.98 | 1.34 (m), 2.06 (m) |
| **8** | 37.17 | - |
| **9** | 37.98 | 1.17 (m), 1.66 (m) |
| **10** | 30.55 | 1.66 (m) |
| **11** | 76.20 | - |
| **12** | 43.68 | - |
| **13** | 26.32 | 1.36 (m), 1.57 (m) |
| **14** | 22.99 | 1.28 (dq, 2.2, 10.7), 1.74 (m) |
| **15** | N.A. | - |
| **16** | 24.36 | 1.14 (s) |
| **17** | 27.24 | 1.06 (s) |
| **18** | 19.22 | 1.03 (s) |
| **19** | 31.38 | 0.90 (d; 0.87) |
| **20** | 20.40 | 1.22 (d; 7.1) |

**Supplementary Table 8. Integrated peak area of oxidized taxanes under different promoters.** Oxidized taxadiene peaks in **Extended Data Fig.6** are integrated by Agilent MassHunter Qualitative Analysis software. The percentage of taxadien-5α-ol (**2**) is highlighted in red.

| | | **35S** | | | | **UBQ10** | | | | **NOS** | | | |
|---|---|---|---|---|---|---|---|---|---|---|---|---|---|
| **oxidized taxadiene** | **name** | rt (min) | area | % | sum % | rt (min) | area | % | sum % | rt (min) | area | % | sum % |
| | iso-OCT (**4**) | 11.3 | 2.7E+04 | 2.7 | | 11.3 | 3.4E+04 | 8.1 | | 11.3 | 1.2E+05 | 9.8 | |
| | OCT (**3**) | 11.9 | 1.3E+05 | 12.6 | | 11.9 | 1.5E+05 | 35.9 | | 11.9 | 4.6E+05 | 37.3 | |
| | | 12.0 | 1.2E+04 | 1.2 | | 12.0 | 5.9E+03 | 1.4 | | 12.0 | 2.8E+04 | 2.3 | |
| **mono** | 5a-ol (**2**) | 12.2 | 2.7E+04 | 2.6 | 31.7 | 12.2 | 3.6E+04 | 8.6 | 70.8 | 12.2 | 1.3E+05 | 10.5 | 75.3 |
| | | 12.3 | 3.2E+04 | 3.1 | | 12.2 | 4.7E+03 | 1.1 | | 12.2 | 1.8E+04 | 1.5 | |
| | **5** | 12.9 | 5.0E+04 | 4.9 | | 12.9 | 5.1E+04 | 12.1 | | 12.9 | 1.6E+05 | 13.4 | |
| | | 13.0 | 3.6E+04 | 3.5 | | 13.0 | 1.5E+04 | 3.6 | | 13.0 | 7.2E+03 | 0.6 | |
| | | 13.2 | 1.0E+04 | 1.0 | | – | – | – | | – | – | – | |
| | | 14.1 | 2.3E+05 | 23.1 | | 13.8 | 1.5E+04 | 3.5 | | 13.9 | 1.6E+04 | 1.3 | |
| | | 14.3 | 1.8E+04 | 1.8 | | 13.9 | 2.2E+04 | 5.3 | | 14.4 | 2.0E+04 | 1.6 | |
| | | 14.5 | 7.9E+04 | 7.8 | | 14.1 | 1.1E+04 | 2.7 | | 14.4 | 3.7E+03 | 0.3 | |
| | | 14.7 | 1.7E+04 | 1.7 | | 14.4 | 3.0E+03 | 0.7 | | 14.5 | 5.2E+03 | 0.4 | |
| | | 14.9 | 6.3E+03 | 0.6 | | 14.5 | 6.9E+03 | 1.6 | | 14.7 | 2.5E+04 | 2.1 | |
| **di** | | 15.1 | 1.4E+04 | 1.4 | 46.1 | 14.8 | 2.2E+03 | 0.5 | 27.4 | 14.8 | 4.8E+03 | 0.4 | 24.7 |
| | | 15.3 | 3.6E+04 | 3.5 | | 15.3 | 2.5E+04 | 5.9 | | 14.9 | 3.6E+03 | 0.3 | |
| | | 15.6 | 2.8E+04 | 2.8 | | 16.4 | 3.0E+04 | 7.2 | | 15.3 | 8.0E+04 | 6.6 | |
| | | 16.2 | 4.1E+03 | 0.4 | | – | – | – | | 16.0 | 2.8E+04 | 2.3 | |
| | | 16.3 | 4.6E+03 | 0.5 | | – | – | – | | 16.2 | 2.8E+04 | 2.3 | |
| | | 16.4 | 2.7E+04 | 2.7 | | – | – | – | | 16.4 | 8.7E+04 | 7.2 | |
| | | 16.6 | 1.6E+05 | 15.5 | | 16.6 | 7.6E+03 | 1.8 | | – | – | – | |
| | | 17.1 | 4.0E+03 | 0.4 | | – | – | – | | – | – | – | |
| **tri** | | 17.3 | 7.8E+03 | 0.8 | 22.2 | – | – | – | 1.8 | – | – | – | 0 |
| | | 17.8 | 1.4E+04 | 1.4 | | – | – | – | | – | – | – | |
| | | 18.1 | 1.2E+04 | 1.2 | | – | – | – | | – | – | – | |
| | | 18.6 | 2.9E+04 | 2.9 | | – | – | – | | – | – | – | |
| **Sum** | | – | 1.0E+06 | 100.0 | 100.0 | – | 4.2E+05 | 100.0 | 100.0 | – | 1.2E+06 | 100.0 | 100.0 |

**Supplementary Table 9. $^{13}C$ & $^{1}H$ δ assignments of 5α,10β-diacetoxy-13α-ol (14) and 5α,10β-diacetoxy-13α-one (15).**

| | 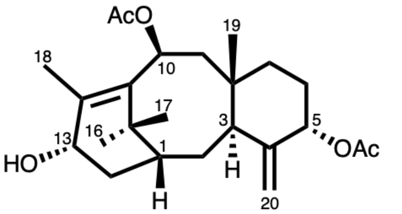<br>taxadien-5α,10β-diacetoxy-13α-ol (**14**) | | 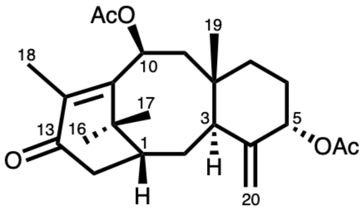<br>taxadien-5α,10β-diacetoxy-13-one (**15**) | |
|---|---|---|---|---|
| **Carbon** | **δ $^{13}C$ (ppm)** | **δ $^{1}H$ (mult.; *J* in Hz)** | **δ $^{13}C$ (ppm)** | **δ $^{1}H$ (mult.; *J* in Hz)** |
| **1** | 40.42 | 1.18 (dd; 4.8, 15.4) | 41.65 | 2.96 (m) |
| **2** | 28.10 | 1.68 (t; 6.2),<br>2.86 (dt; 15.2, 9.7) | 26.92 | 1.94 (d; 19.6),<br>2.91 (dd; 7.3, 19.6) |
| **3** | 37.11 | 3.04 (m) | 36.11 | 2.15 (m) |
| **4** | N.A. | - | N.A. | - |
| **5** | 76.96 | 5.34 (s) | 75.75 | 5.29 (s) |
| **6** | 28.25 | 1.79 (m), 1.86 (m) | 28.18 | 1.79 (m) |
| **7** | 34.10 | 1.25 (m), 2.02 (m) | 33.28 | 1.27 (m), 1.99 (m) |
| **8** | 38.48 | - | 38.45 | - |
| **9** | 44.22 | 1.59 (dd; 5.5, 14.8),<br>2.32 (t; 13.4) | 43.53 | 1.70 (dd; 5.5, 14.5),<br>2.43 (t; 13.4) |
| **10** | 71.39 | 6.11 (dd; 5.5, 12.3) | 71.66 | 6.13 (dd; 5.6, 12.2) |
| **11** | 137.27 | - | 155.30 | - |
| **12** | 137.91 | - | 136.10 | - |
| **13** | 68.40 | 4.41 (m) | 200.47 | - |
| **14** | 36.78 | 1.73 (d; 9.6) | 36.11 | 1.28 (m), 1.79 (m) |
| **15** | 38.98 | - | 39.92 | - |
| **16** | 32.47 | 0.95 (s) | 37.31 | 1.13 (s) |
| **17** | 25.90 | 1.48 (s) | 25.05 | 1.57 (s) |
| **18** | 22.02 | 2.18 (s) | 21.55 | 1.95 (s) |
| **19** | 21.63 | 0.72 (s) | 22.13 | 0.75 (s) |
| **20** | 113.26 | 4.83 (s), 5.15 (s) | 112.45 | 4.78 (s), 5.14 (s) |

| | | | | |
|---|---|---|---|---|
| **-<u>C</u>$O_2$Me** | 170.36 | - | 169.85 | - |
| **-<u>C</u>$O_2$Me** | 170.36 | - | 170.04 | - |
| **-$CO_2$<u>Me</u>** | 21.50 | 2.05 (s) | 21.43 | 2.09 (s) |
| **-$CO_2$<u>Me</u>** | 15.95 | 2.10 (s) | 13.66 | 2.18 (s) |

* N.A.:  chemical shift not assigned due to insufficient information

**Supplementary Table 10. Accession numbers of genes used in this study.**

| Name | Organism | Accession number |
|---|---|---|
| TS1 | *Taxus brevefolia* | U48796 |
| TS2 | *Taxus canadensis* | AY364470 |
| T5αH | *Taxus cuspidata* | AY289209 |
| TAT | *Taxus cuspidata* | AF190130 |
| T10βH | *Taxus cuspidata* | AF318211 |
| DBAT | *Taxus baccata* | AF456342 |
| T13αH | *Taxus cuspidata* | AY056019 |

**Supplementary Table 11. List of PCR primer pairs used in this study.**

| Purpose | Target | | Sequence (5' to 3') |
|---|---|---|---|
| Cloning | T5αH | F | ATTCTGCCCAAATTCGCGACCGGT |
| | | R | GAAACCAGAGTTAAAGGCCTCGAGCTATGGTCTCGGAAACAGTTTAAT |
| | TAT | F | ATTCTGCCCAAATTCGCGACCGGT |
| | | R | GAAACCAGAGTTAAAGGCCTCGAGTCATACTTTAGCCACATATTTTTT |
| | T10βH | F | ATTCTGCCCAAATTCGCGACCGGT |
| | | R | GAAACCAGAGTTAAAGGCCTCGAGTTAGGATCTCGGAAAAAGTTTTAT |
| | DBAT | F | ATTCTGCCCAAATTCGCGACCGGT |
| | | R | GAAACCAGAGTTAAAGGCCTCGAGTCAAGGTTTAGTTACATATTTGTT |
| | T13αH | F | ATTCTGCCCAAATTCGCGACCGGT |
| | | R | GAAACCAGAGTTAAAGGCCTCGAGTTAAGATCTGGAATAGAGTTTAAT |
| Promoter switching | To replace 35S with NOS in the pEAQ | F | CAATTAGAGTCTCATATTCACTCTCAATTATTAAAATCTTAATAGGTTTTG ATAAAAGCG |
| | | R | CTAAAGAAAATTTAATGAAACCAGAGTTAAACCGGTCGCGAATTTGGG CAGAATATACAG |
| | To replace 35S with UBQ10 in the pEAQ | F | ATTAATCTGAGTTTTTCTGATTAACACTTATTAAAATCTTAATAGGTTTTG ATAAAAGCG |
| | | R | CTAAAGAAAATTTAATGAAACCAGAGTTAAACCGGTCGCGAATTTGGG CAGAATATACAG |
| | To PCR any gene from pEAQ vector into either pEAQ-NOS or pEAQ-UBQ10 | F | CAATTAGAGTCTCATATTCACTCTCAATTATTAAAATCTTAATAGGTTTTG ATAAAAGCG |
| | | R | GTAAATTCAAACTAAAGAAAATTTAATGAAACCAGAGTTAA |

Nucleotides underlined are overlaps designed for Gibson assembly into pEAQ vectors. Synthetic genes contain 5'-overlaps designed for Gibson assembly into pEAQ vectors thus the same forward primer is used for PCR.

**Supplementary Table 12. List of *S. cerevisiae* strains used in this study.**

| Strain ID | Name/alias | Description | Genotype | Parent strain |
|---|---|---|---|---|
| JBEI-18127 | TS-expressing/ 5xTS | Mevalonate pathway, 3x CrtE GGPP synthases, 5x TS with protein tags, all in galactose-inducible promoters. | CEN.PK2-1C {1114a,1622b,308a,911b::GAL1p-MBP-TXS-ERG20-TDH1t; 1014a::GAL1p-TXS-GFP-ADH1t, leu2-3, 112::HIS3MX6-GAL1p-ERG19/GAL10p-ERG8; ura3-52::ura3/GAL1p-MvaSA110G/GAL10p-MvaE; his3Δ1::hphMX4-GAL1p-ERG12/GAL10p-IDI1; trp1-289::TRP1/GAL1p-CrtE/GAL10p-ERG20; YPRCdelta15::NatMX-GAL1p-CrtE/GAL10p-CrtE; MATa} | JWy1 |
| JBEI-18128 | TS+T5αH-expressing/ 3xTS {T5αH-CPR} | Mevalonate pathway, 3x CrtE GGPP synthases, 3x TS with protein tags, and 1x T5αH-CPR, all in galactose-inducible promoters. | CEN.PK2-1C {511b::GAL1p-T5OH-PGK1t/GAL3p-CPR-ENO2t; 1114a,1622b::GAL1p-MBP-TXS-ERG20-TDH1t; 1014a::GAL1p-TXS-GFP-ADH1t, leu2-3, 112::HIS3MX6-GAL1p-ERG19/GAL10p-ERG8; ura3-52::ura3/GAL1p-MvaSA110G/GAL10p-MvaE; his3Δ1::hphMX4-GAL1p-ERG12/GAL10p-IDI1; trp1-289::TRP1/GAL1p-CrtE/GAL10p-ERG20; YPRCdelta15::NatMX-GAL1p-CrtE/GAL10p-CrtE; MATa} | JWy1 |

Methods used to construct these strains are previously described.[13]

### Supplementary Figures

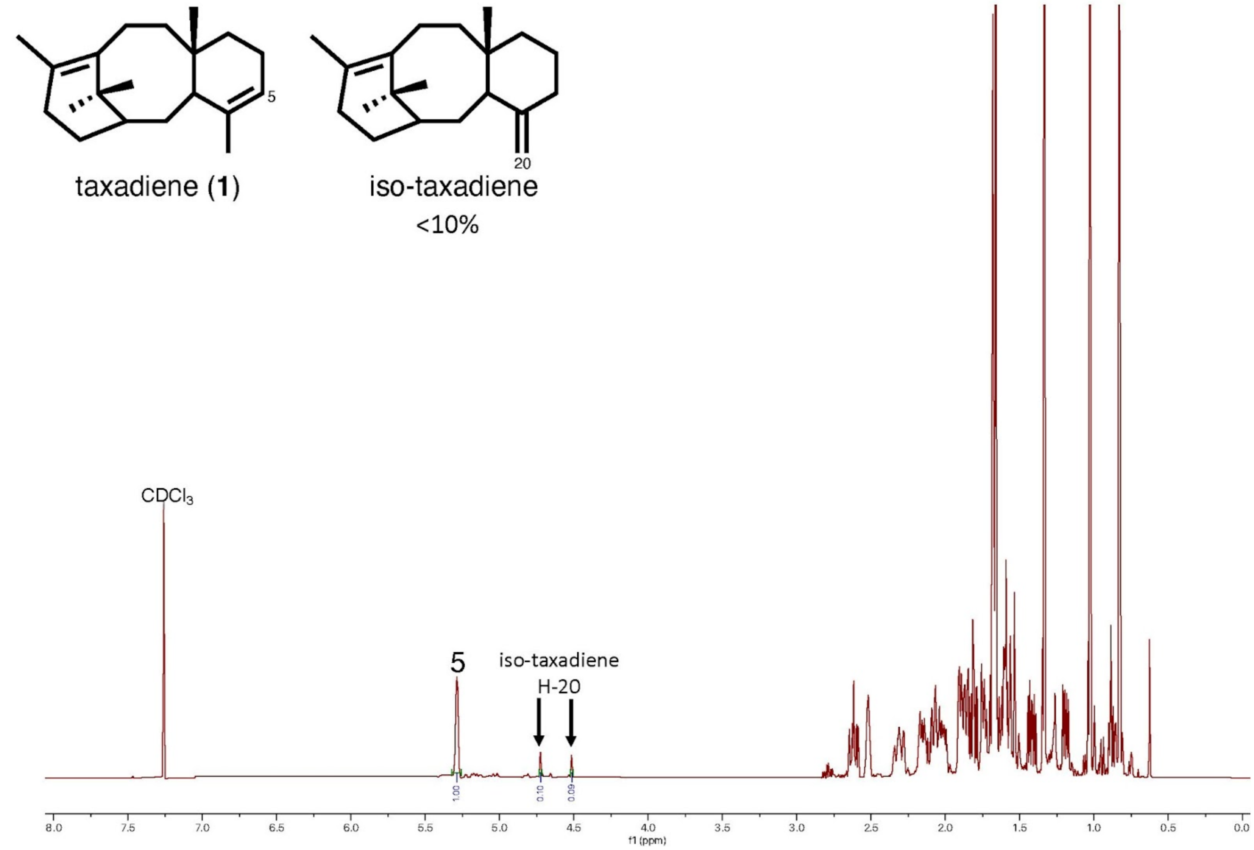


**Supplementary Figure 1. $^1$H-NMR spectrum of taxadiene (1) in $CDCl_3$ (600 Hz, n = 16).**

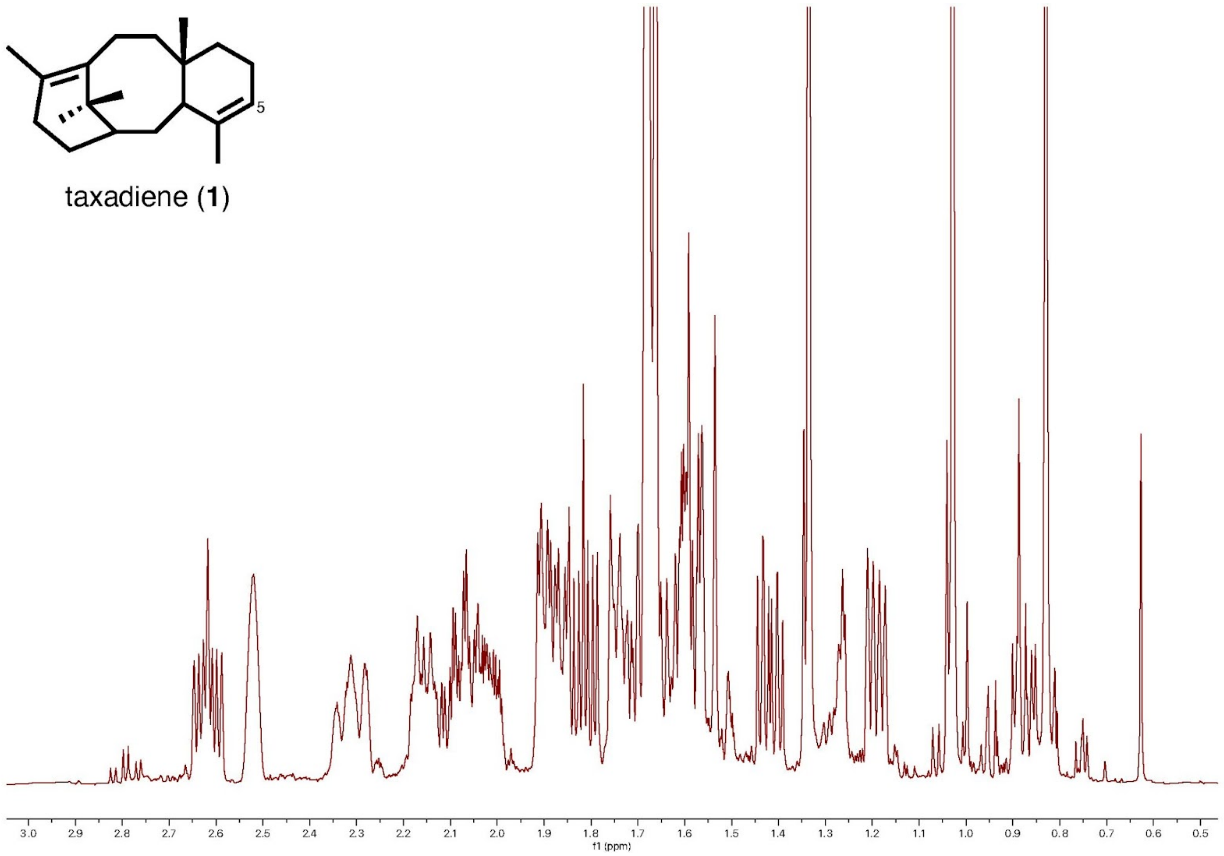


**Supplementary Figure 2. $^1$H NMR spectrum of taxadiene (1) in $CDCl_3$ (600 Hz, n = 16) in the region of 0.5~3.0 ppm.**

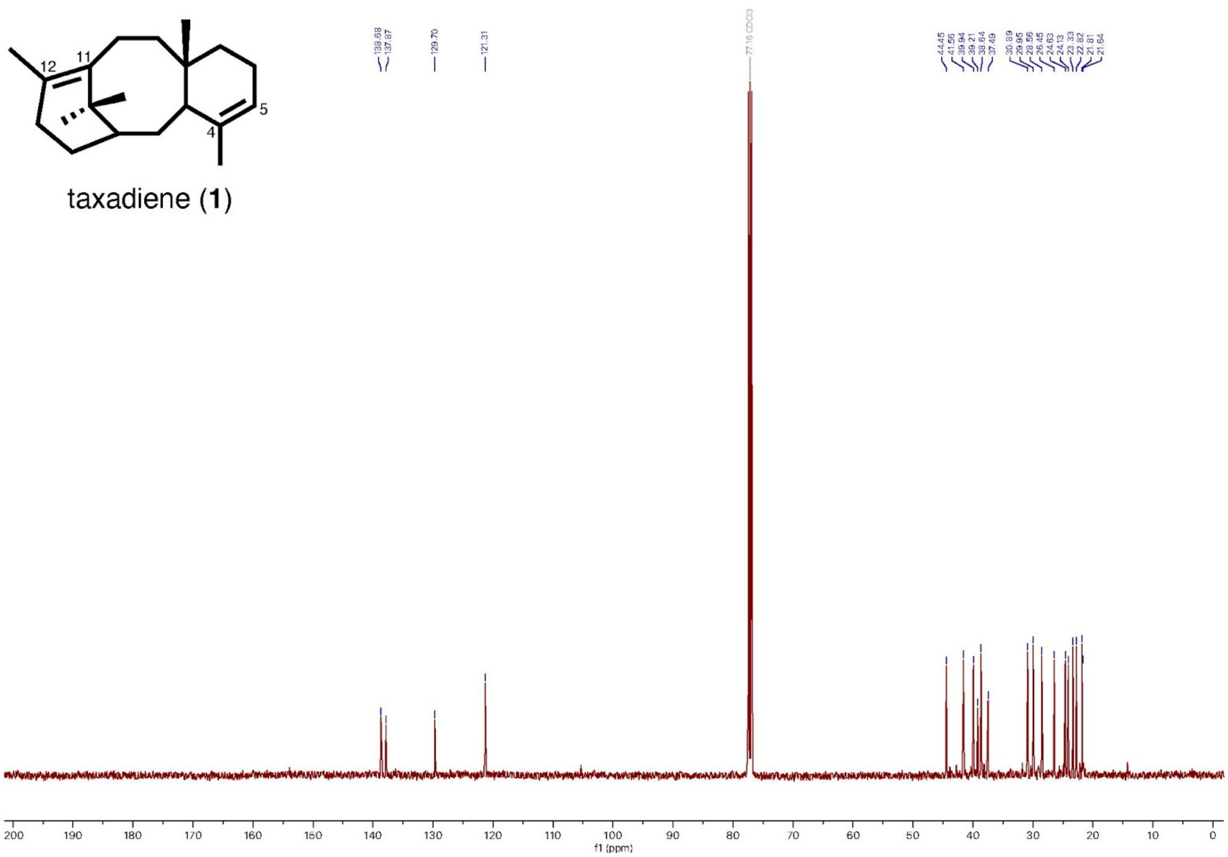


**Supplementary Figure 3. $^{13}$C NMR spectrum of taxadiene (1) in $CDCl_3$ (500 Hz, n = 512).** Chemical shifts (ppm): 138.68, 137.87, 129.70, 121.31, 44.45, 41.56, 39.94, 39.21, 38.64, 37.49, 30.89, 29.95, 28.56, 26.45, 24.63, 24.13, 23.33, 22.82, 21.81, 21.64.

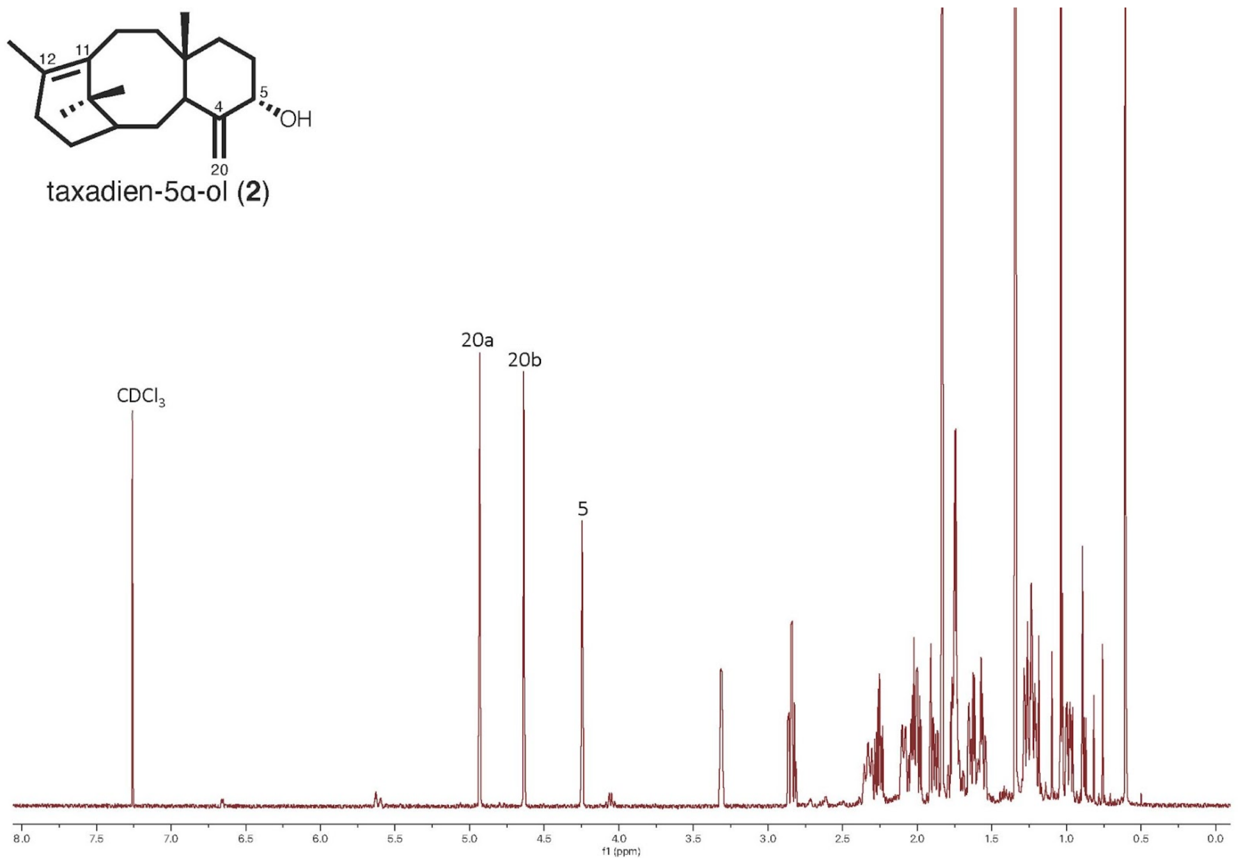


**Supplementary Figure 4. $^1$H-NMR spectrum of taxadien-5α-ol (2) in $CDCl_3$ (600 Hz, n = 32).**

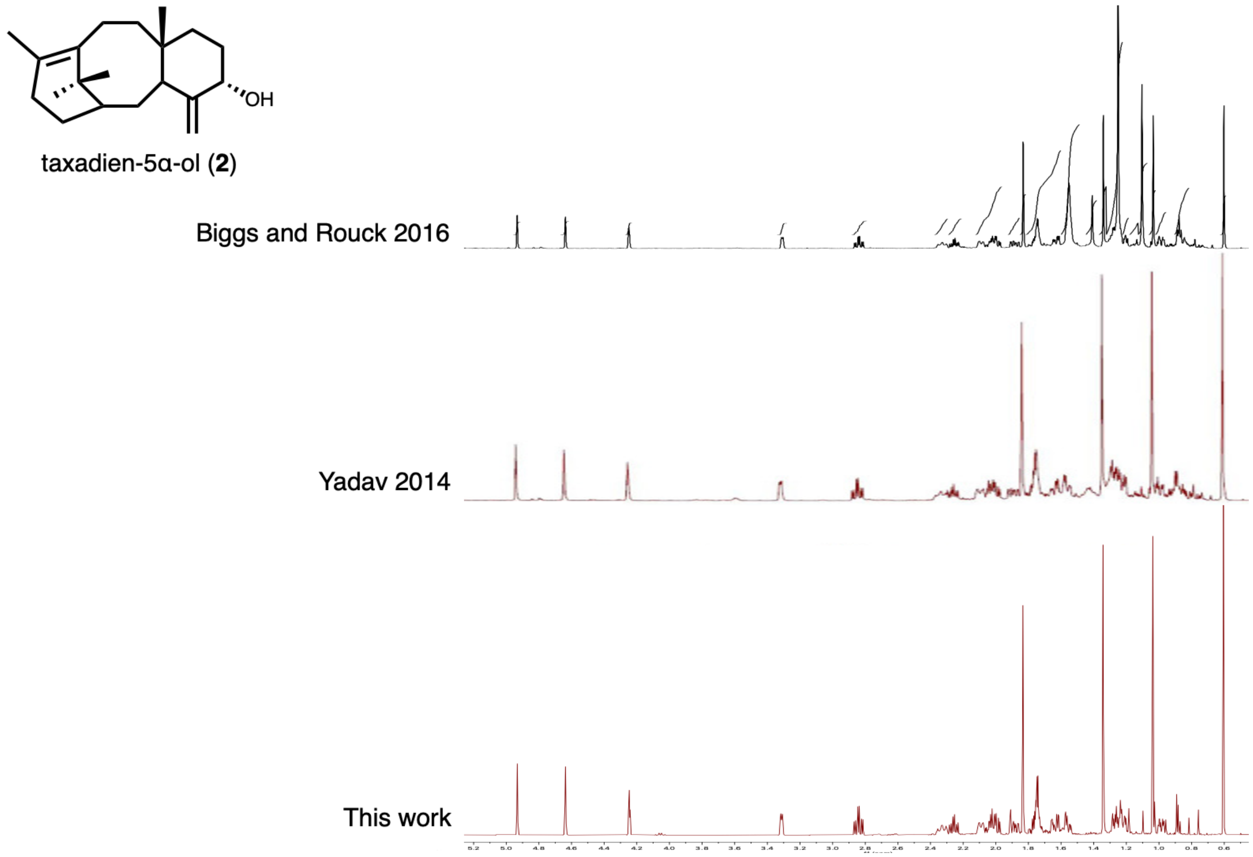


**Supplementary Figure 5. $^{1}$H-NMR spectra of taxadien-5α-ol (2) from previous works and this work in the region of 0.6~5.2 ppm.**

1H-NMR spectrum of **2** purified from engineered *E. coli* expressing TS and T5αH in two previous works[4,6] (top 2 traces) are used to confirm the identity of the synthetic **2** in this work.

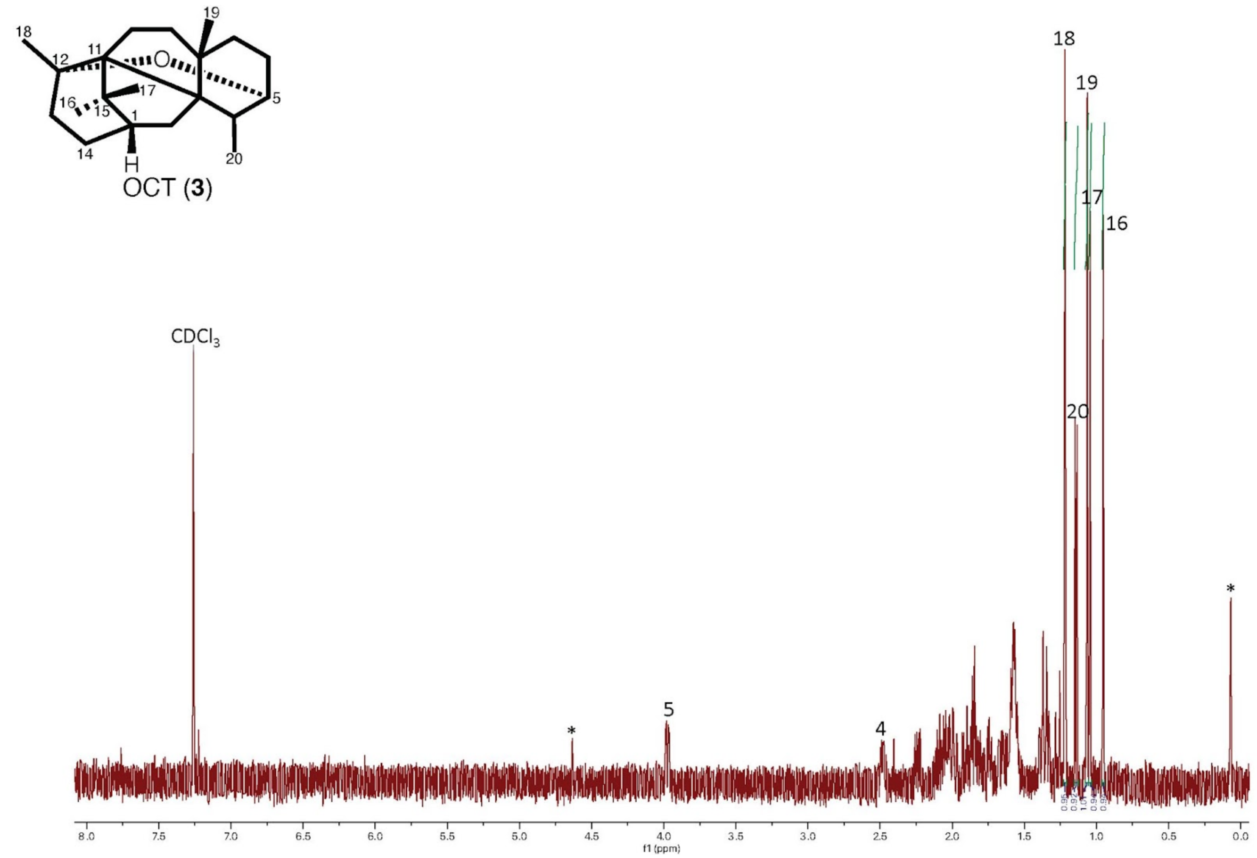


**Supplementary Figure 6. $^1$H-NMR spectrum of OCT (3) in $CDCl_3$ (500 Hz, n = 64).** Asterisk indicates impurities.

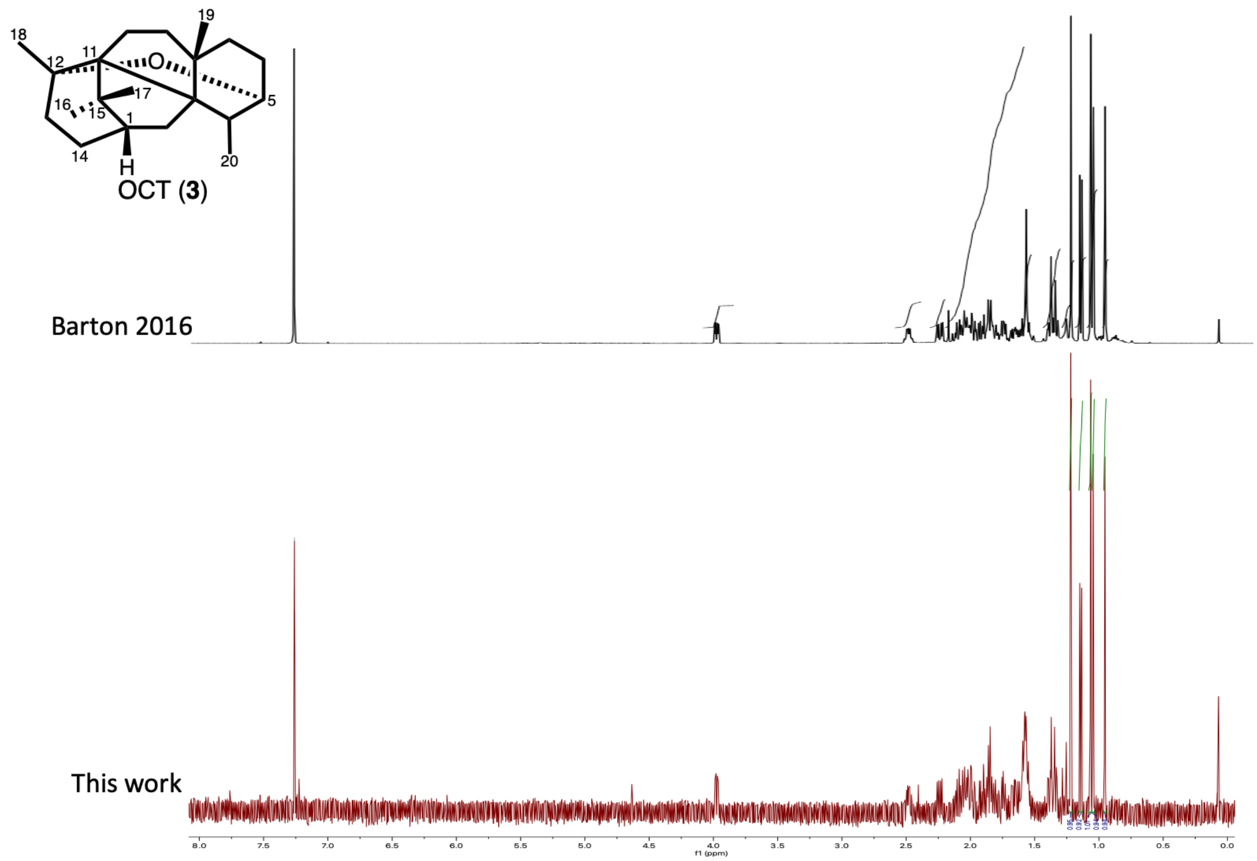


**Supplementary Figure 7. $^{1}$H-NMR spectra of OCT (3) from previous work and this work.** $^{1}$H-NMR spectra of synthetic OCT[15] (top trace) and OCT (**3**) isolated from tobacco in this work (bottom trace).

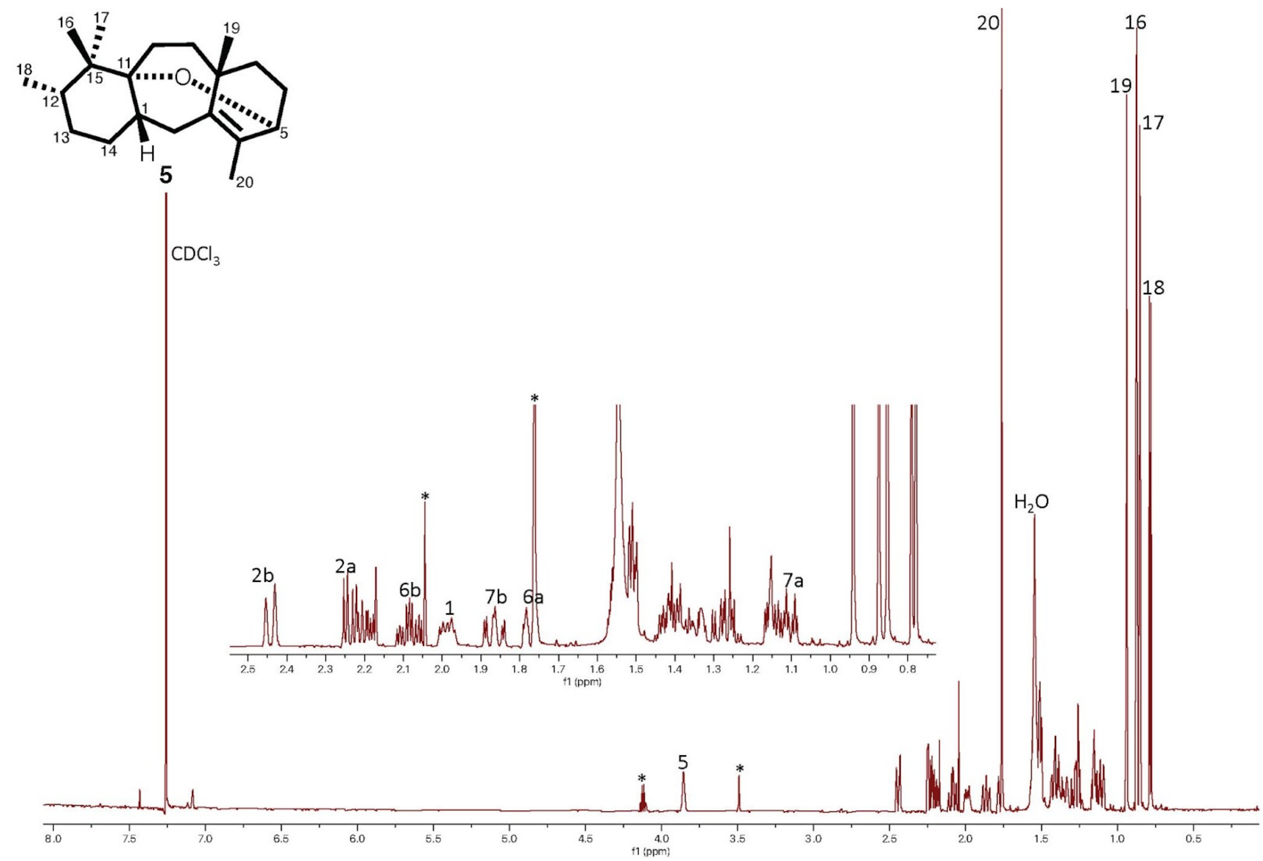


**Supplementary Figure 8. $^{1}$H-NMR spectrum of compound 5 in $CDCl_3$ (600 Hz, n = 32).** Asterisk indicates impurities.

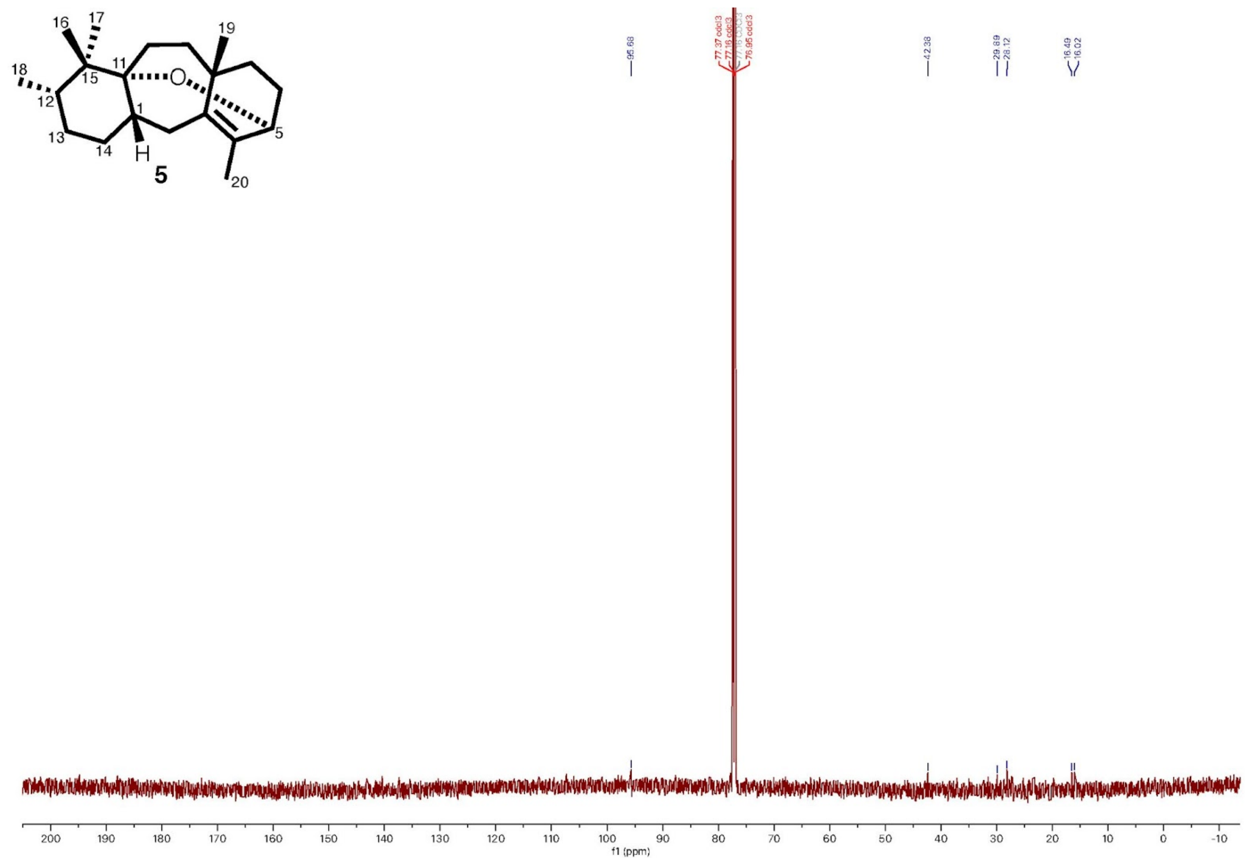


**Supplementary Figure 9. $^{13}$C-NMR spectrum of compound 5 in $CDCl_3$ (600 Hz, n = 2048).**

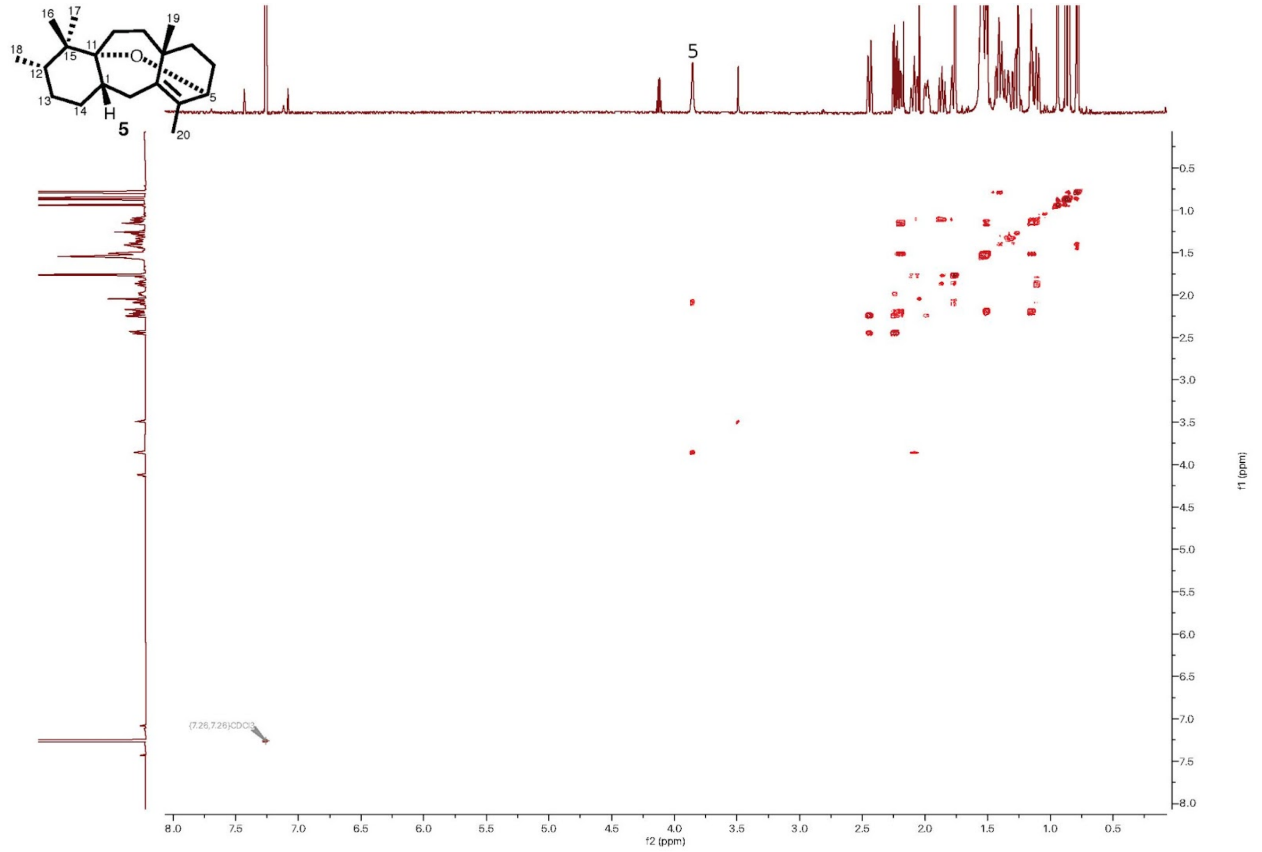


**Supplementary Figure 10. COSY spectrum of compound 5 in $CDCl_3$ (600 Hz, n = 4).**

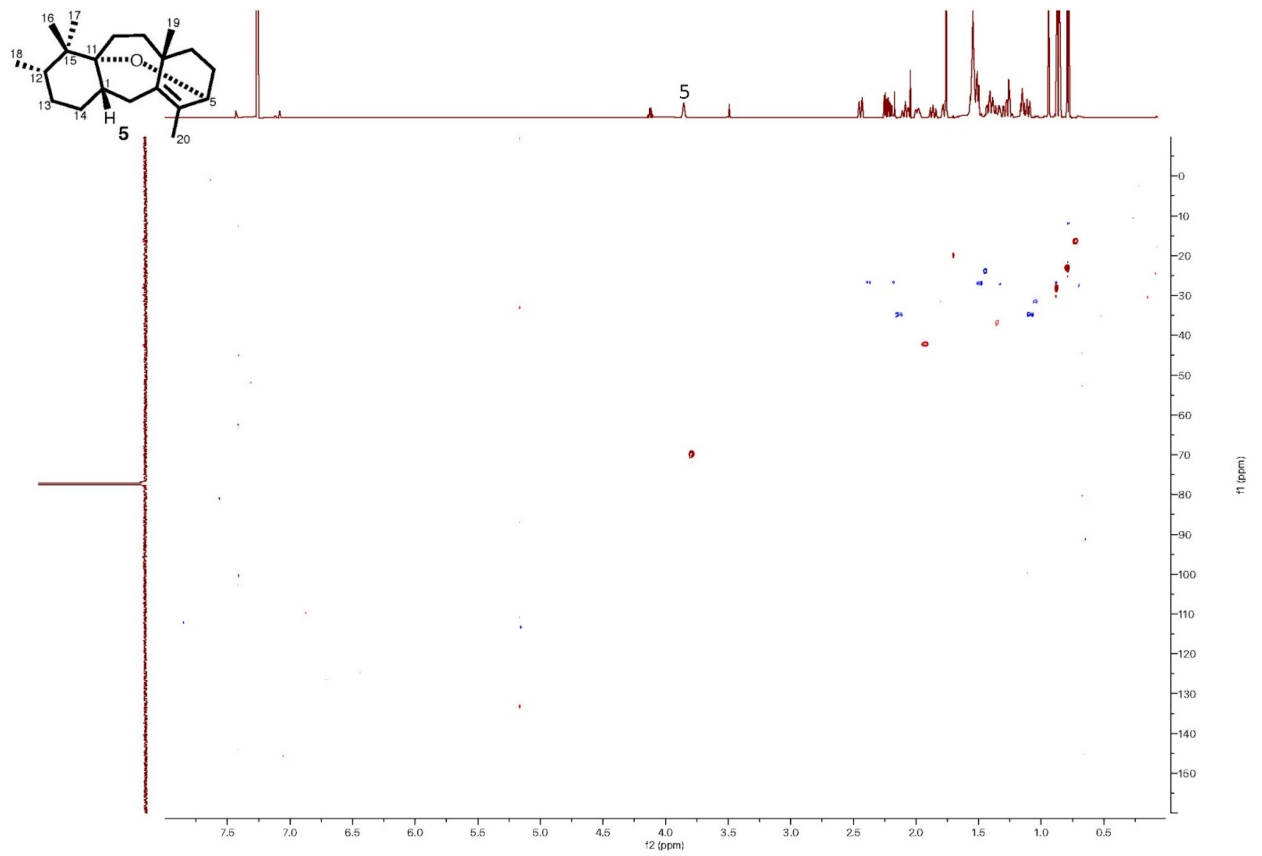


**Supplementary Figure 11. HSQC spectrum of compound 5 in $CDCl_3$ (600 Hz, n = 8).**

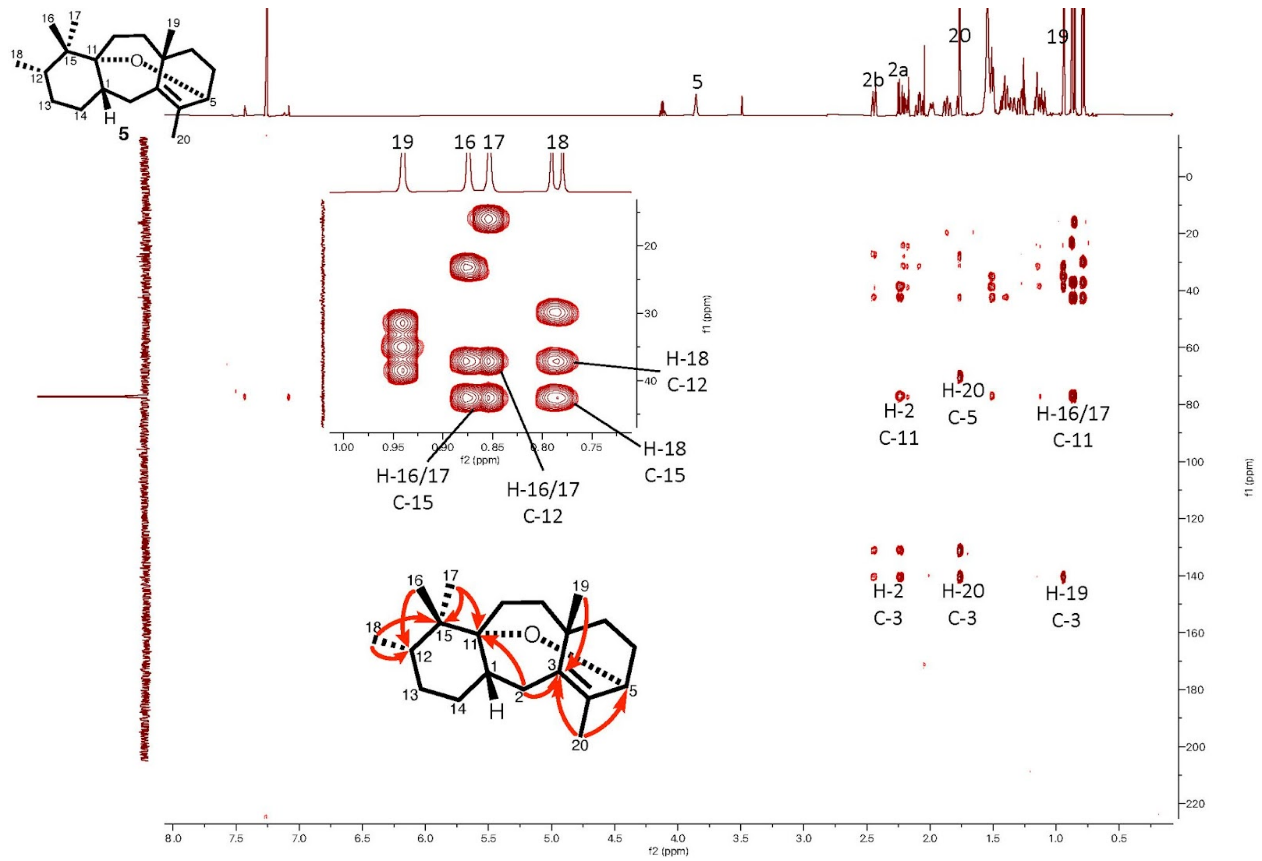


**Supplementary Figure 12. HMBC spectrum of compound 5 in $CDCl_3$ (600 Hz, n = 32).**

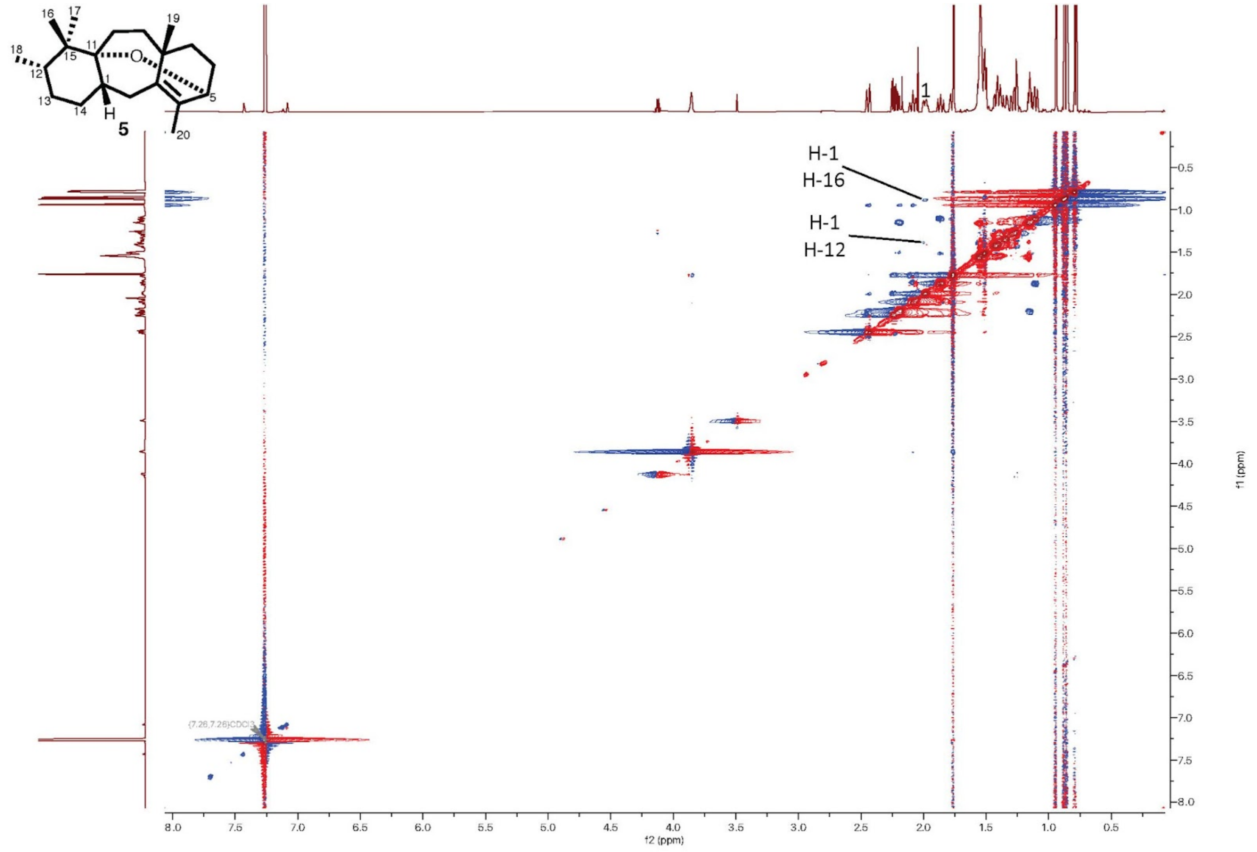


**Supplementary Figure 13. ROESY spectrum of compound 5 in CDCl$_3$ (600 Hz, n = 8).**

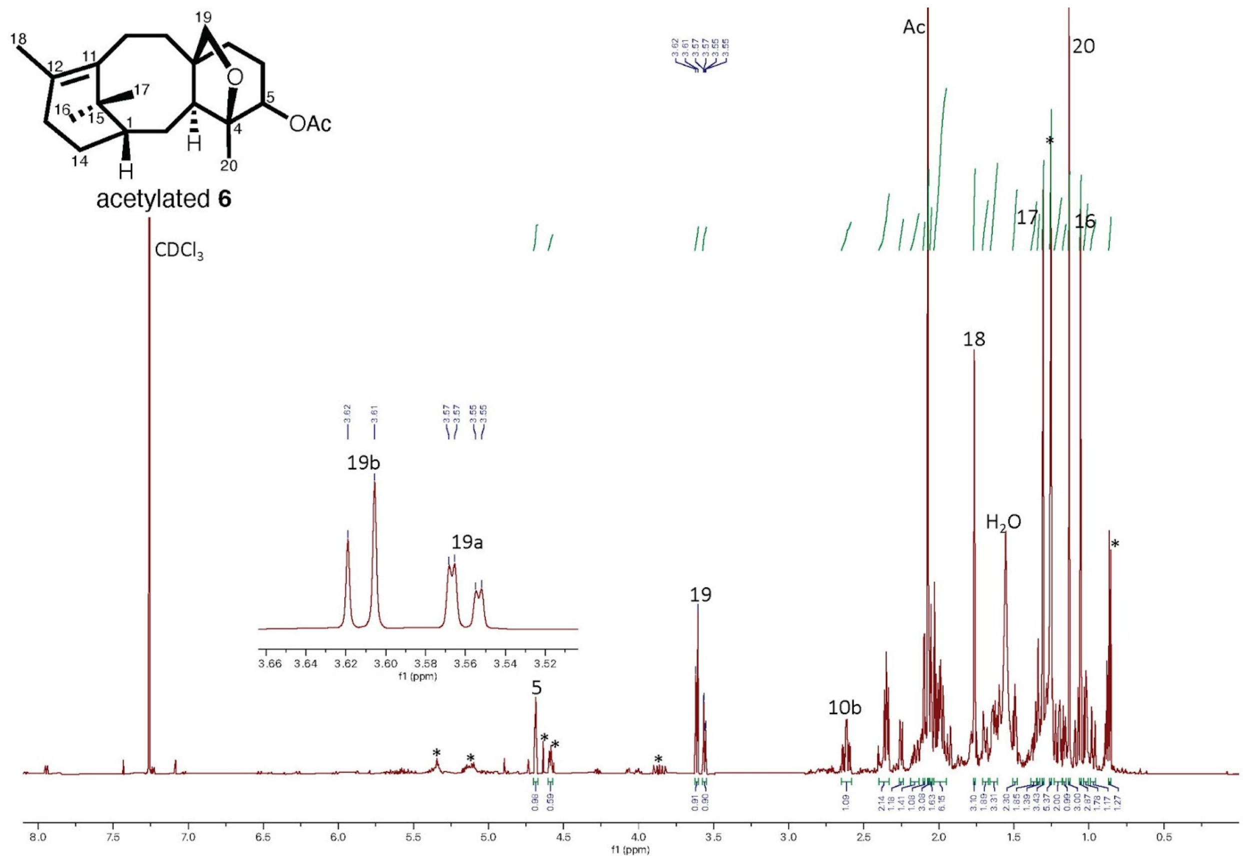


**Supplementary Figure 14. $^{1}$H-NMR spectrum of acetylated 6 in $CDCl_3$ (600 Hz, n = 64).** Asterisk indicates impurities. The W-coupling of proton 19a is shown.

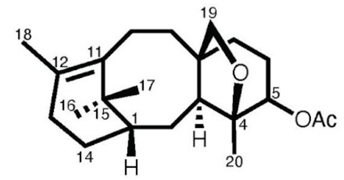


acetylated **6**

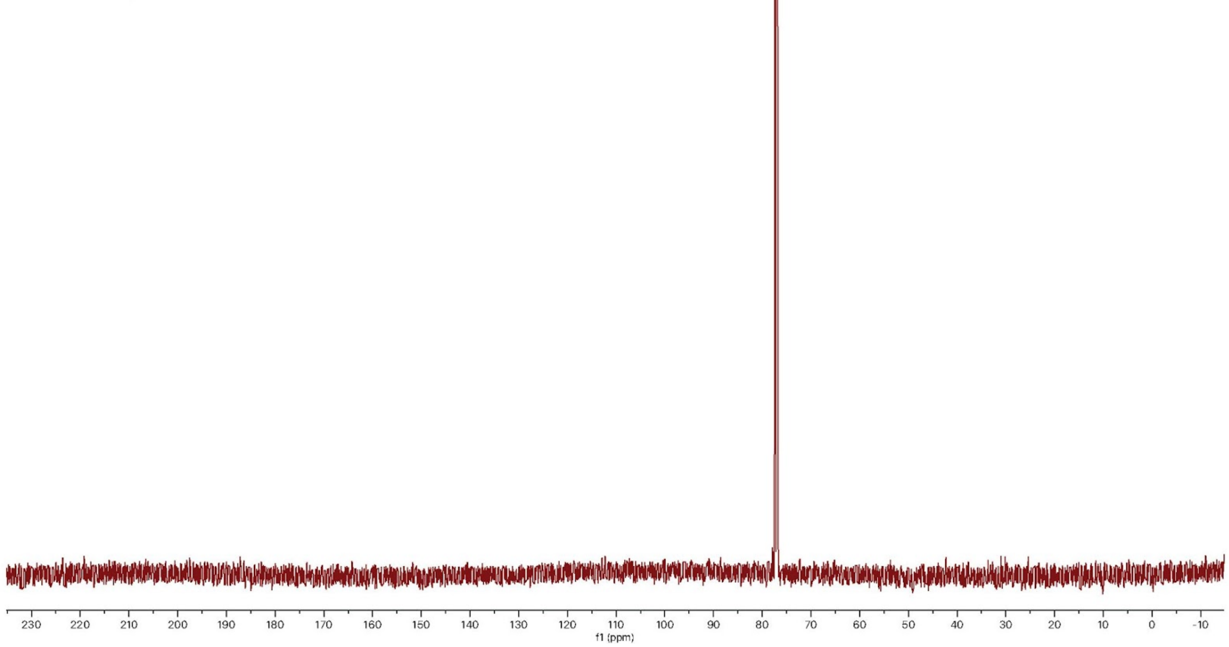


**Supplementary Figure 15. $^{13}$C-NMR spectrum of acetylated 6 in $CDCl_3$ (600 Hz, n = 1024).**

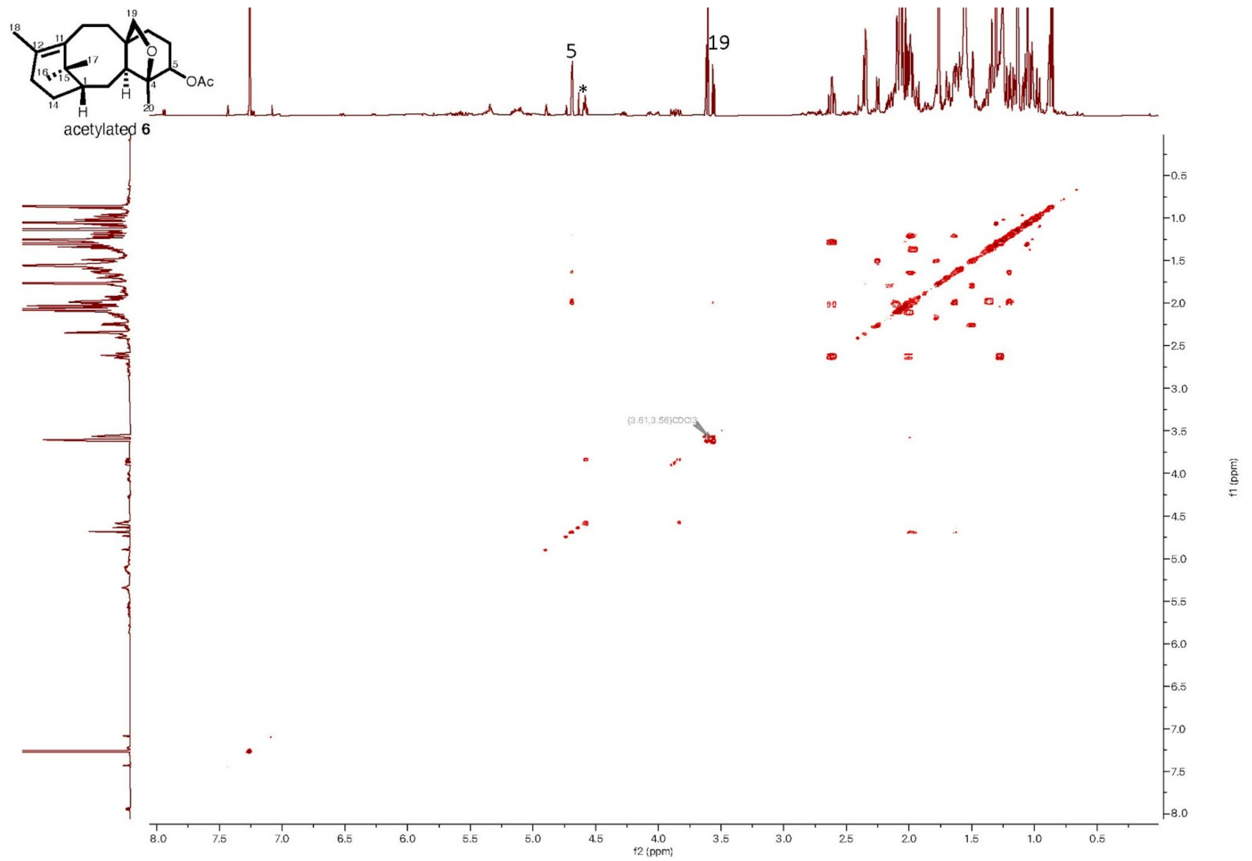


**Supplementary Figure 16. COSY spectrum of acetylated 6 in $CDCl_3$ (600 Hz, n = 4).**

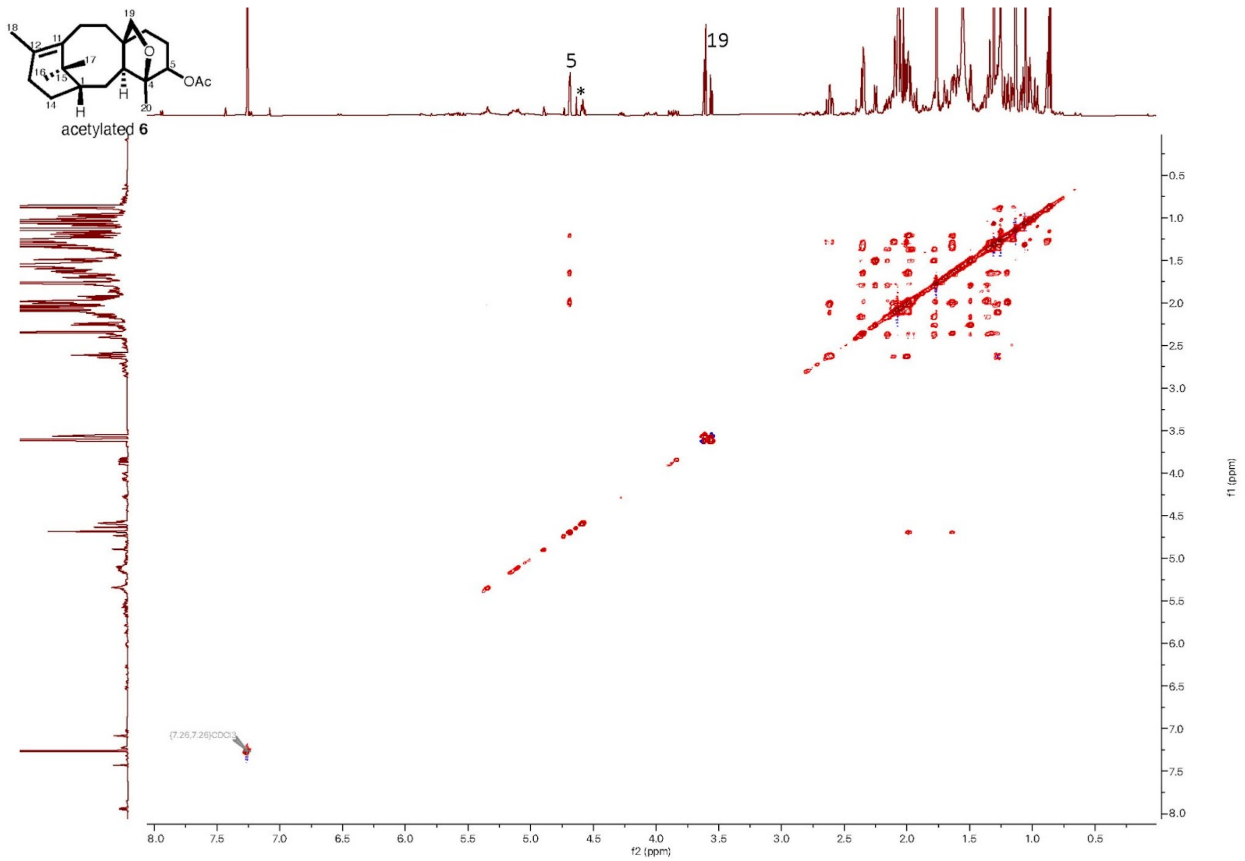


**Supplementary Figure 17. TOCSY spectrum of acetylated 6 in $CDCl_3$ (600 Hz, n = 2).**

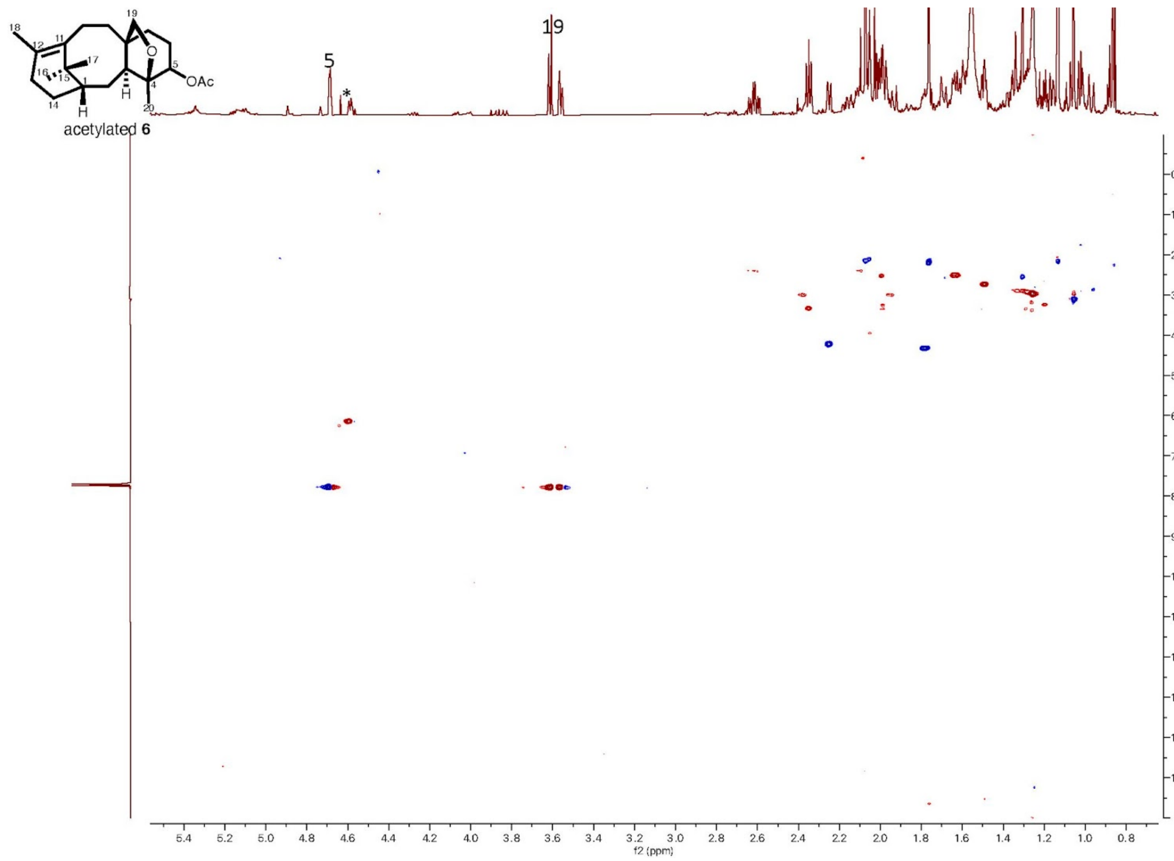


**Supplementary Figure 18. HSQC spectrum of acetylated 6 in $CDCl_3$ (600 Hz, n = 8).**

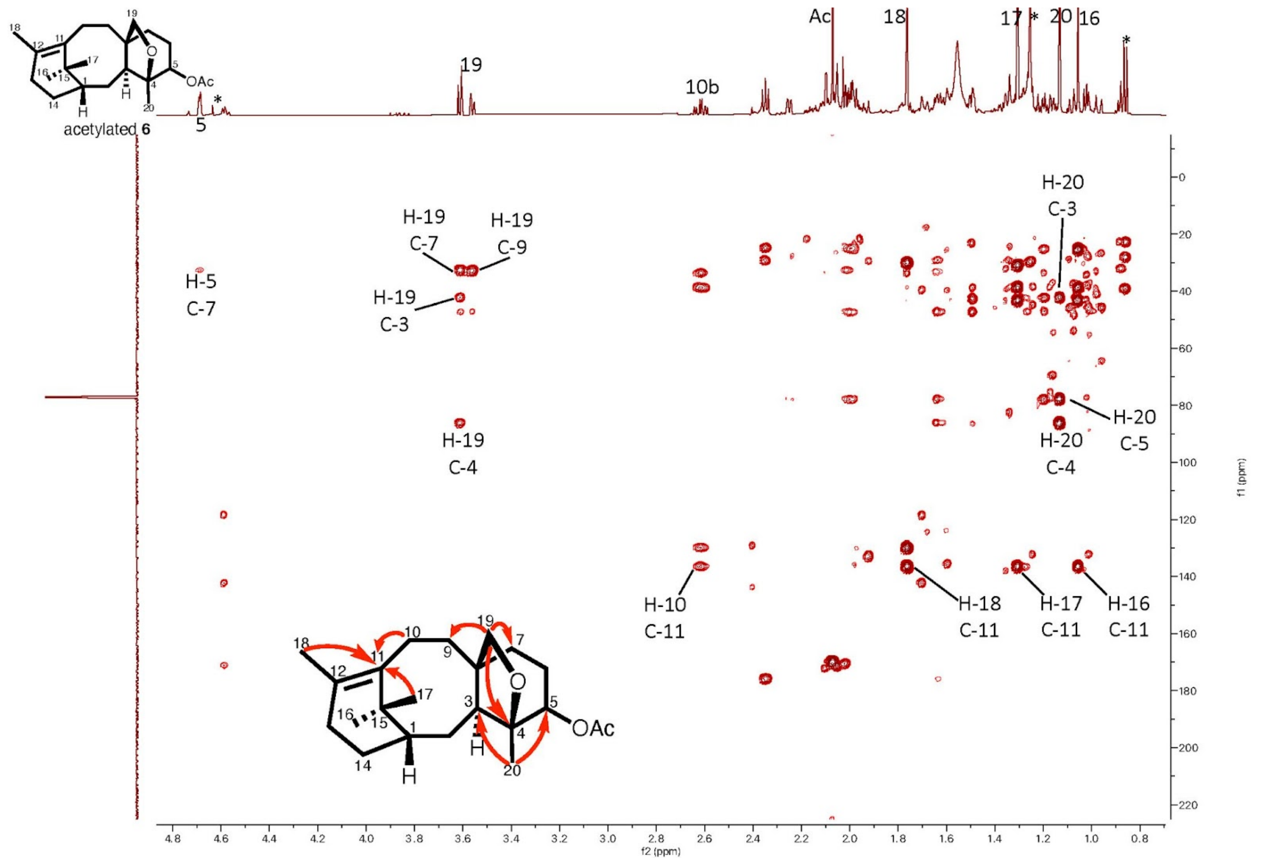


**Supplementary Figure 19. HMBC spectrum of acetylated 6 in $CDCl_3$ (600 Hz, n = 32).**

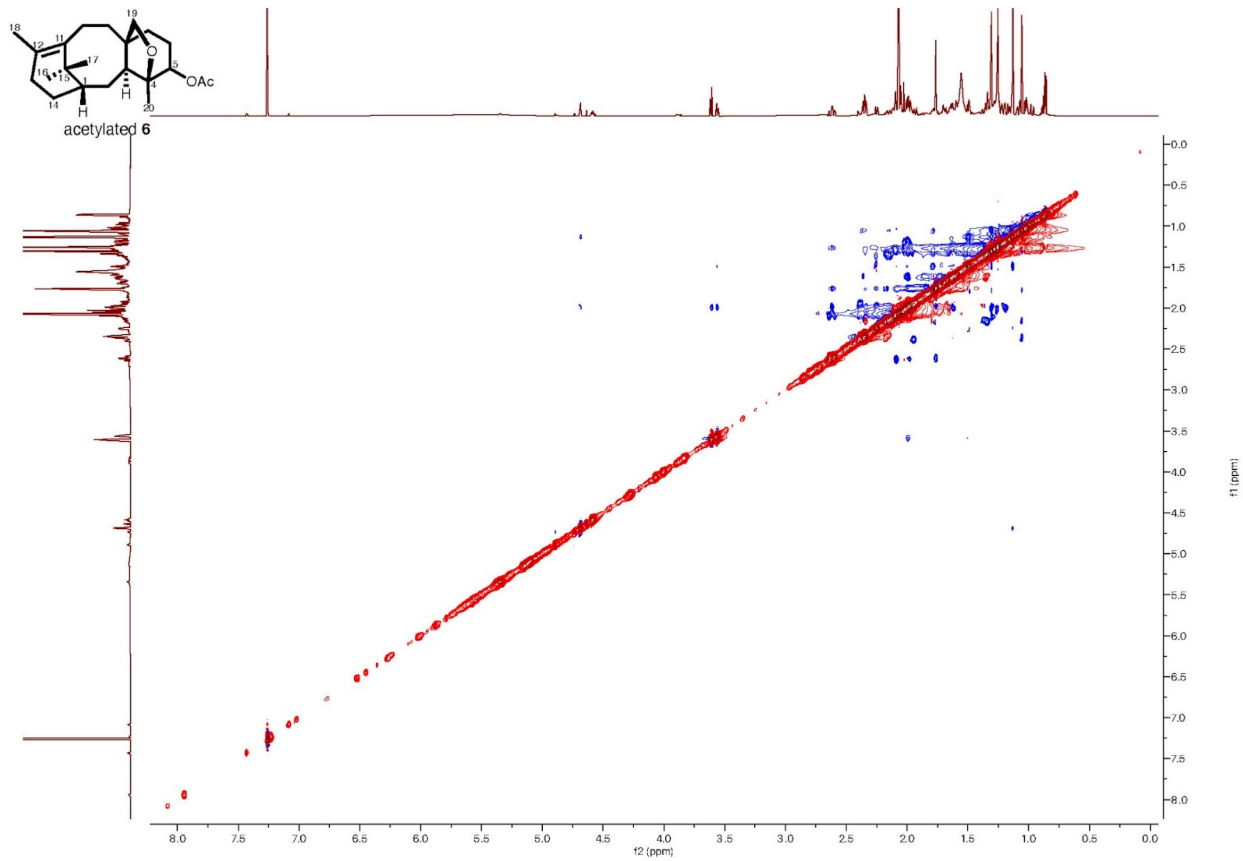


**Supplementary Figure 20. ROESY spectrum of acetylated 6 in $CDCl_3$ (600 Hz, n = 8).**

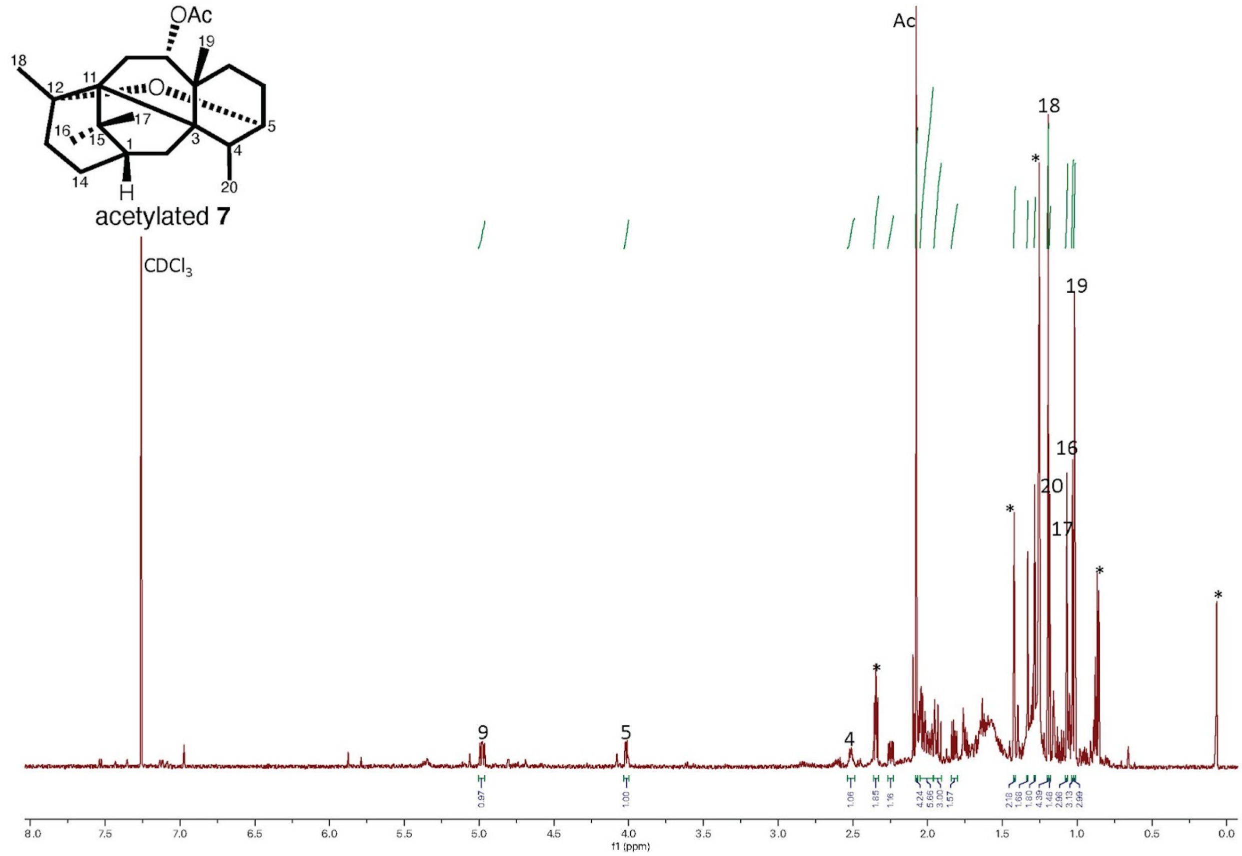


**Supplementary Figure 21. $^1$H-NMR spectrum of acetylated 7 in $CDCl_3$ (600 Hz, n = 64).** Asterisk indicates impurities.

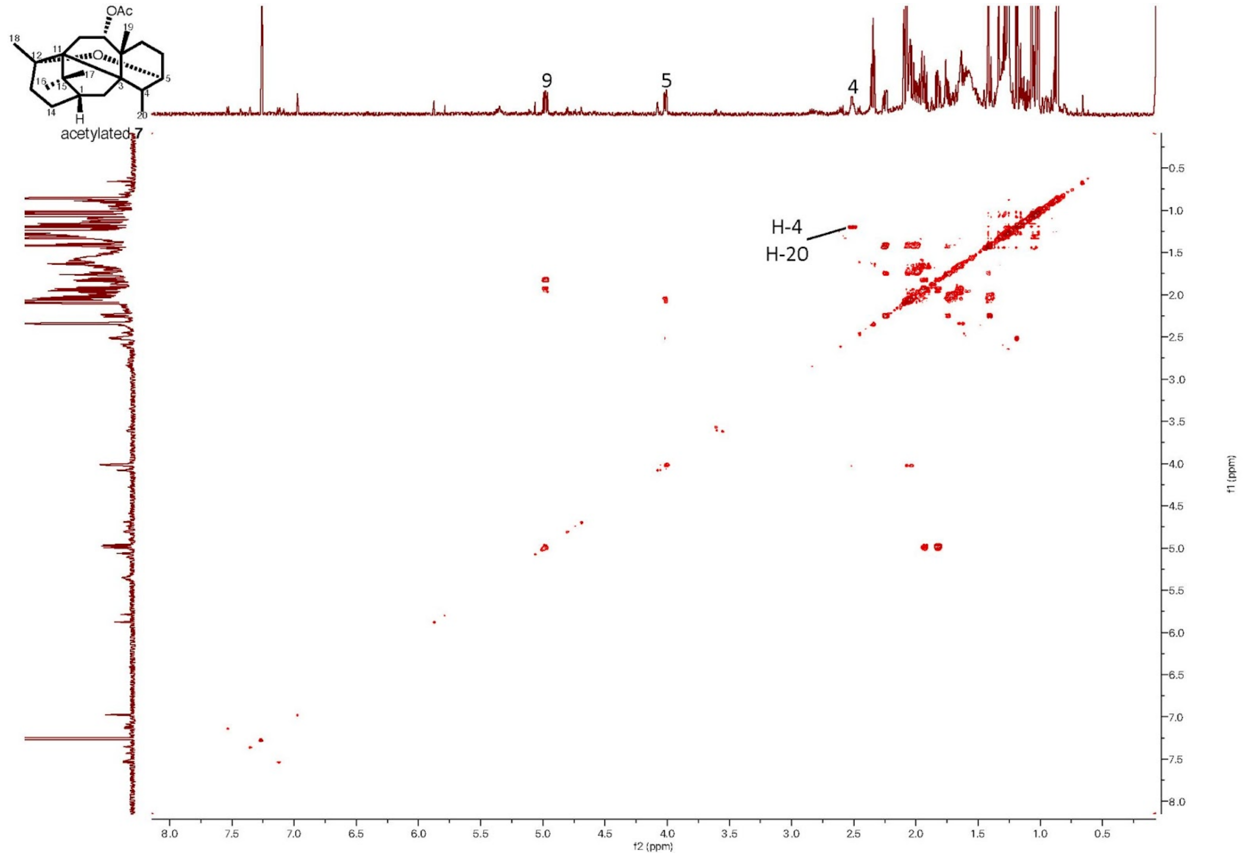


**Supplementary Figure 22. COSY spectrum of acetylated 7 in $CDCl_3$ (600 Hz, n = 4).**

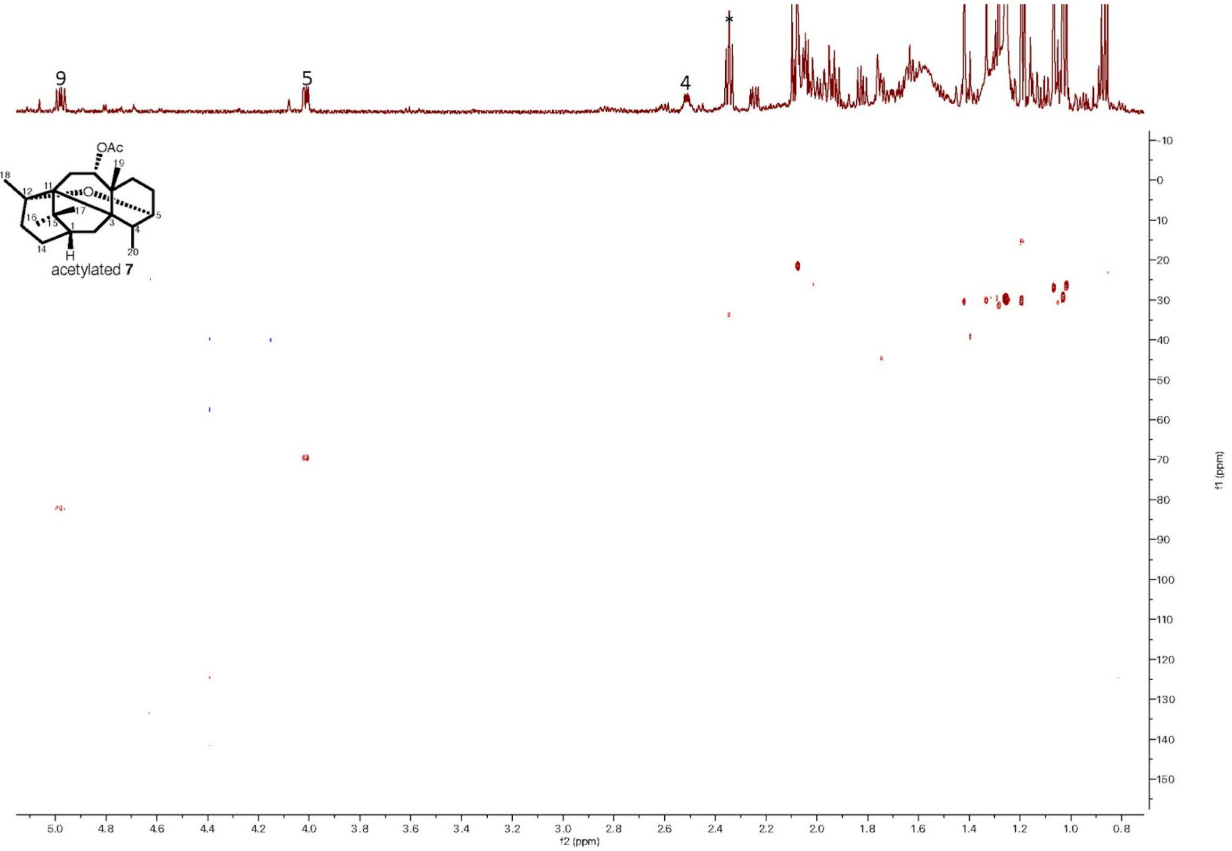


**Supplementary Figure 23. HSQC spectrum of acetylated 7 in $CDCl_3$ (600 Hz, n = 8).**

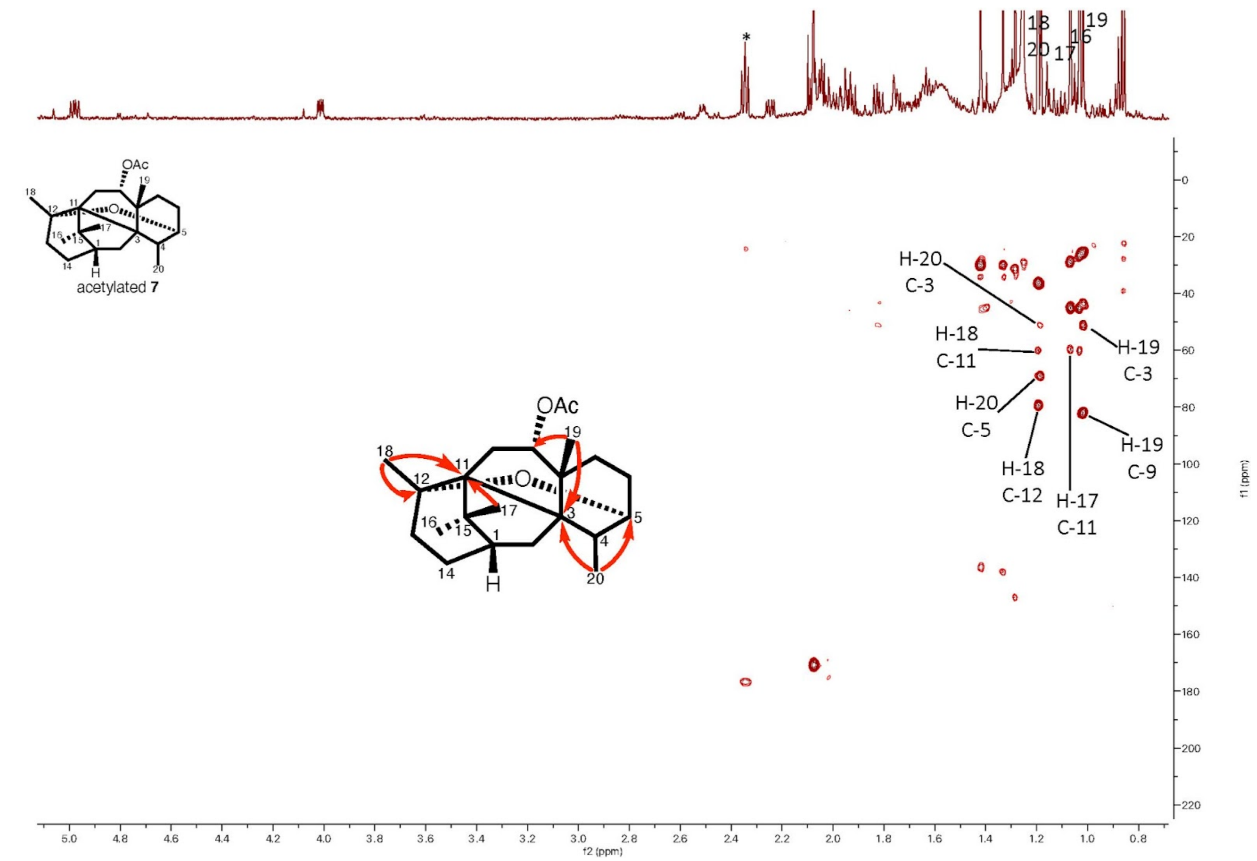


**Supplementary Figure 24. HMBC spectrum of acetylated 7 in $CDCl_3$ (600 Hz, n = 32).**

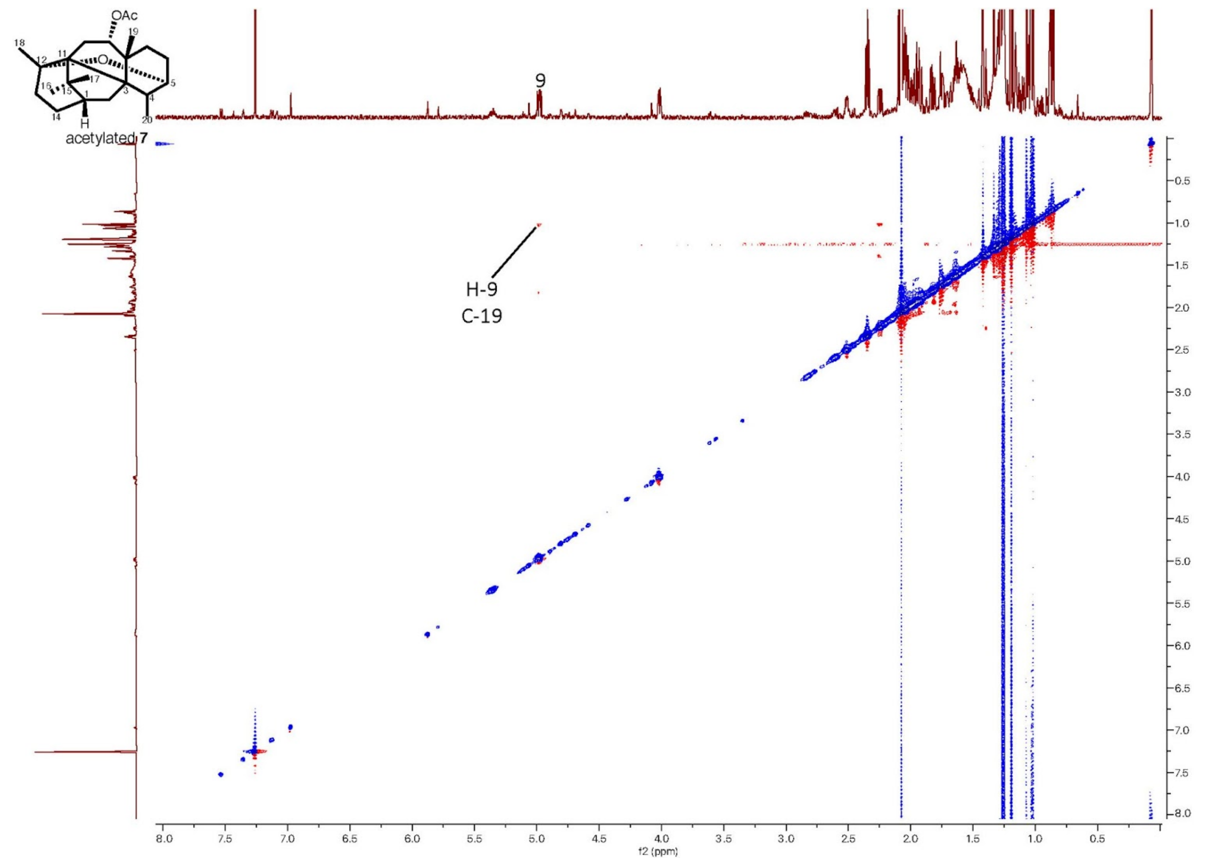


**Supplementary Figure 25. ROESY spectrum of acetylated 7 in $CDCl_3$ (600 Hz, n = 8).**

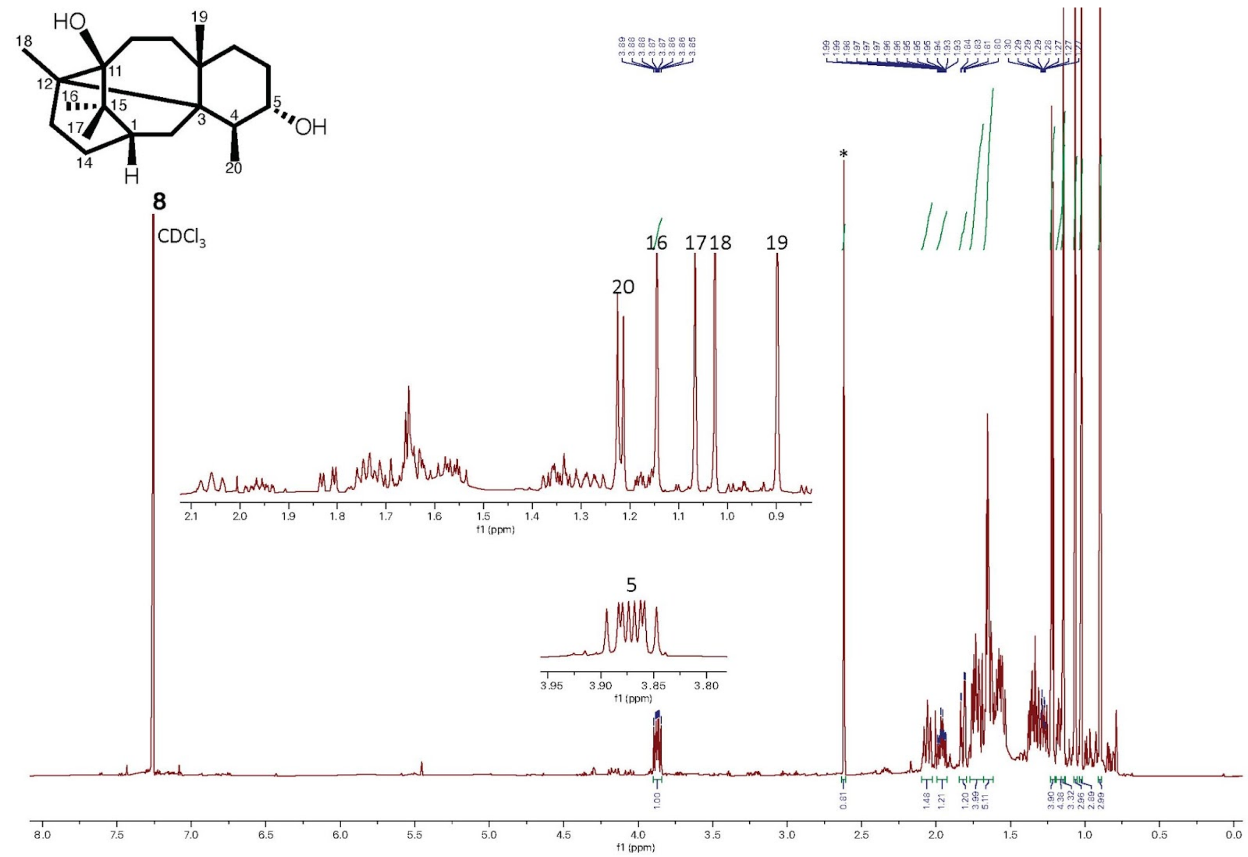


**Supplementary Figure 26. $^{1}$H-NMR spectrum of compound 8 in $CDCl_3$ (600 Hz, n = 64).**
Asterisk indicates impurities.

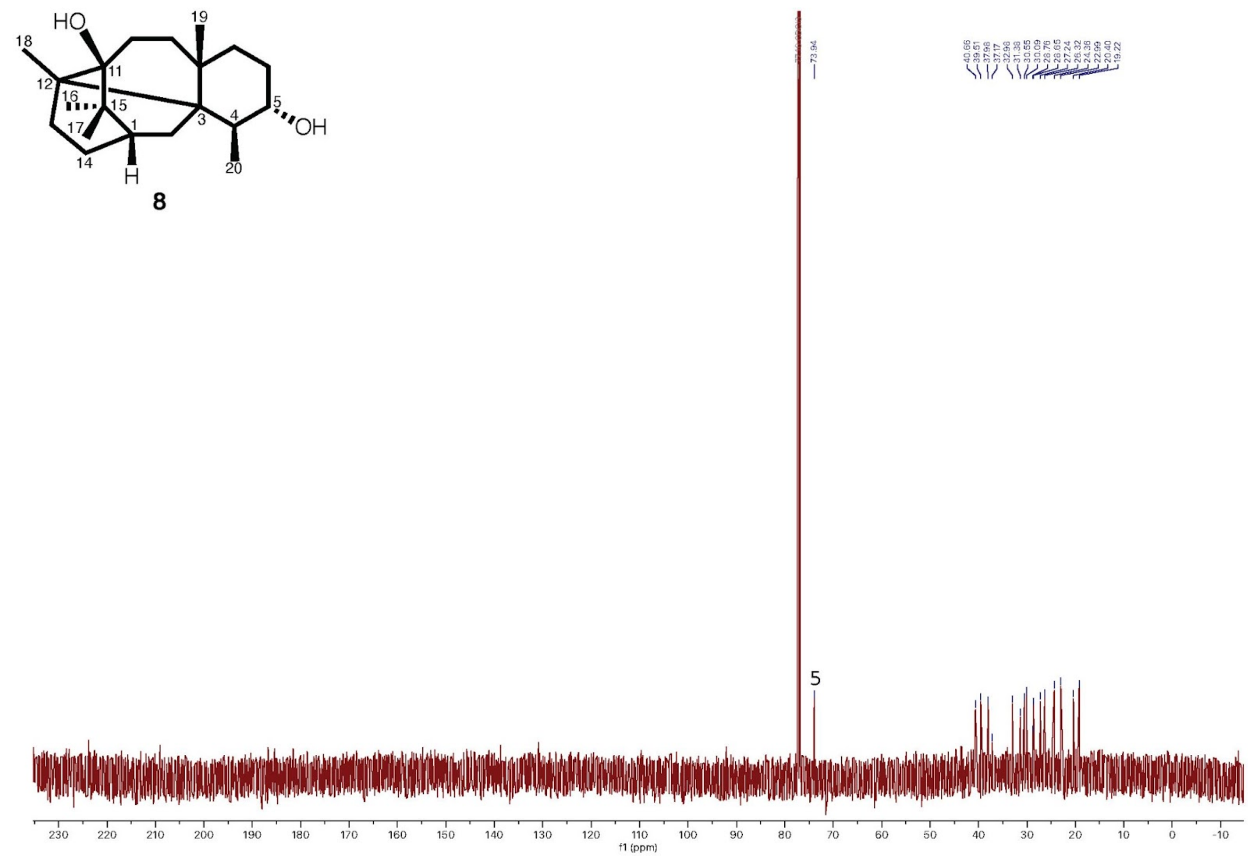


**Supplementary Figure 27. $^{13}$C-NMR spectrum of compound 8 in $CDCl_3$ (600 Hz, n = 1024).**

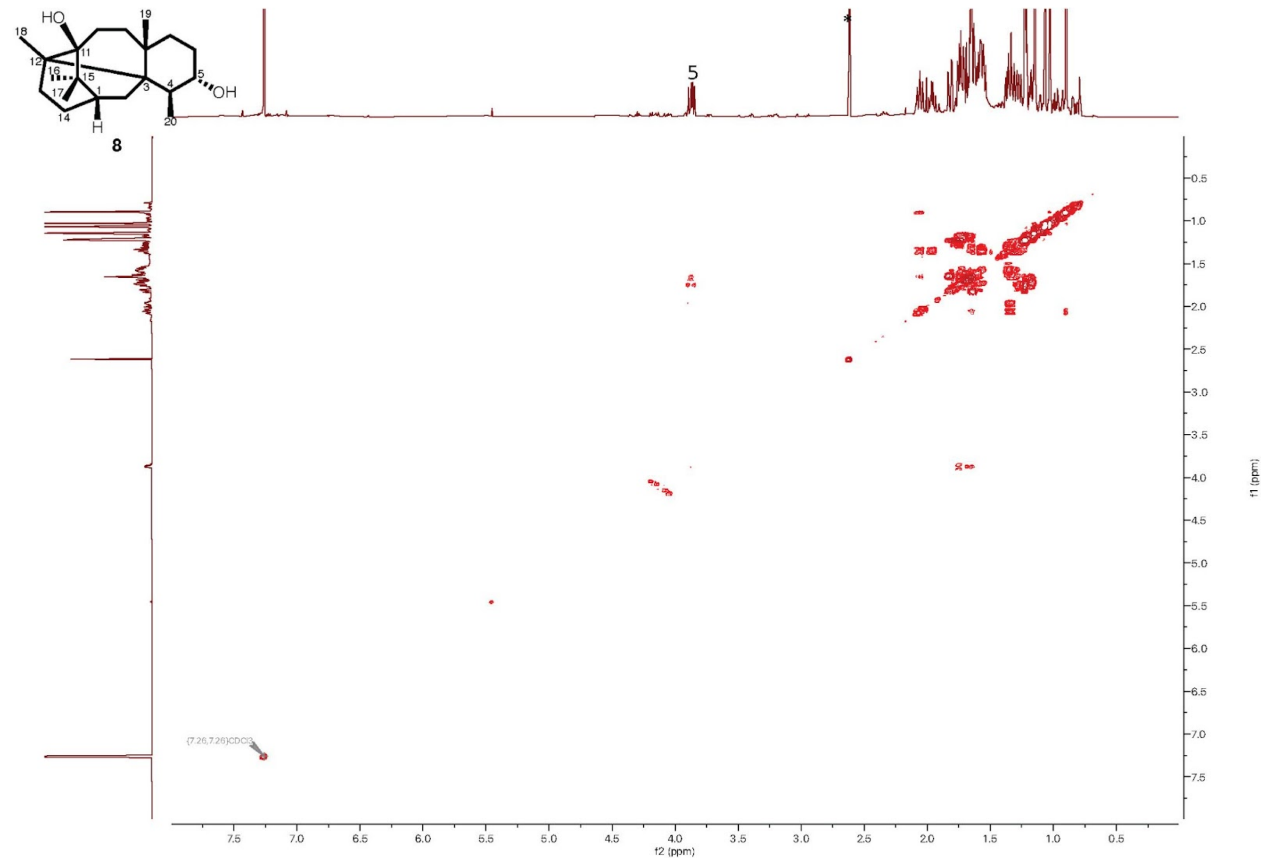


**Supplementary Figure 28. COSY spectrum of compound 8 in $CDCl_3$ (600 Hz, n = 8).**

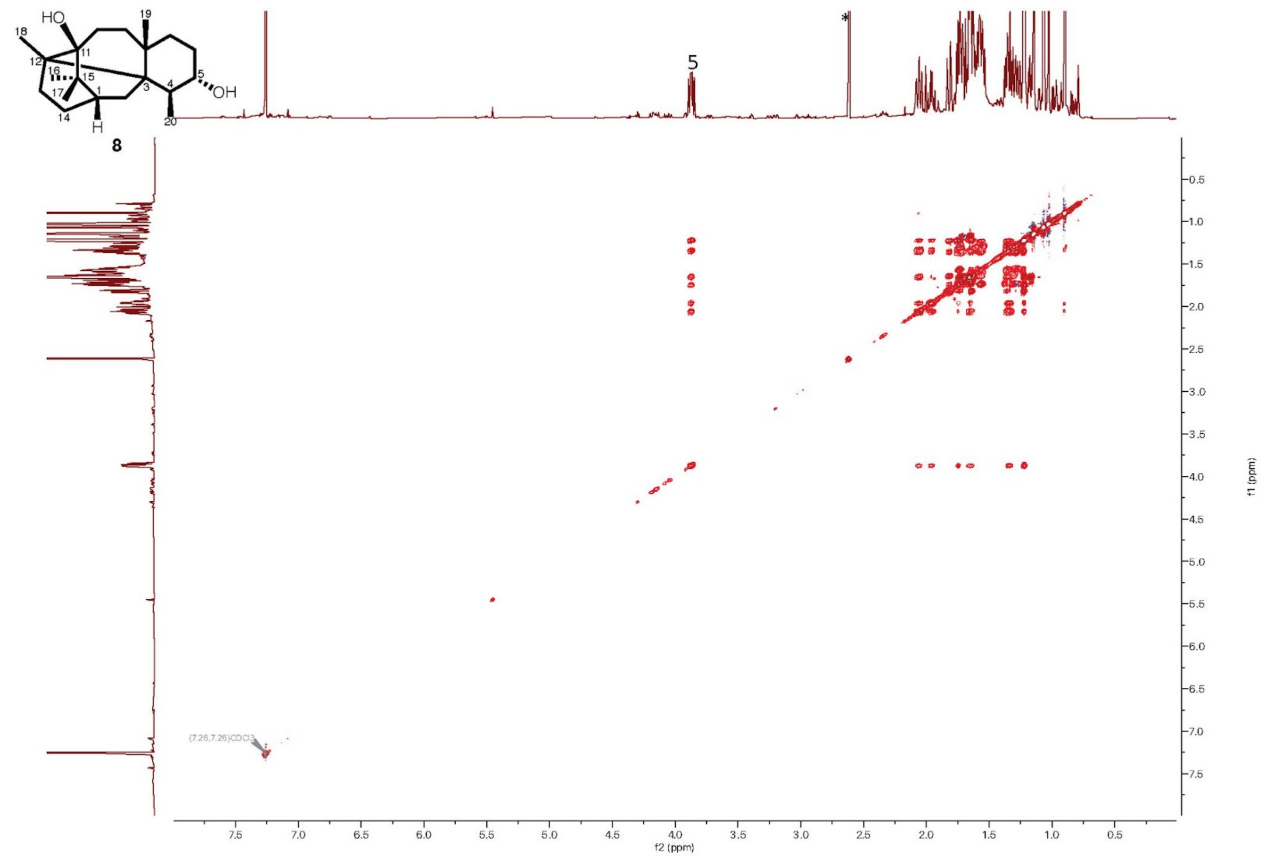

**Supplementary Figure 29. TOCSY spectrum of compound 8 in $CDCl_3$ (600 Hz, n = 4).**

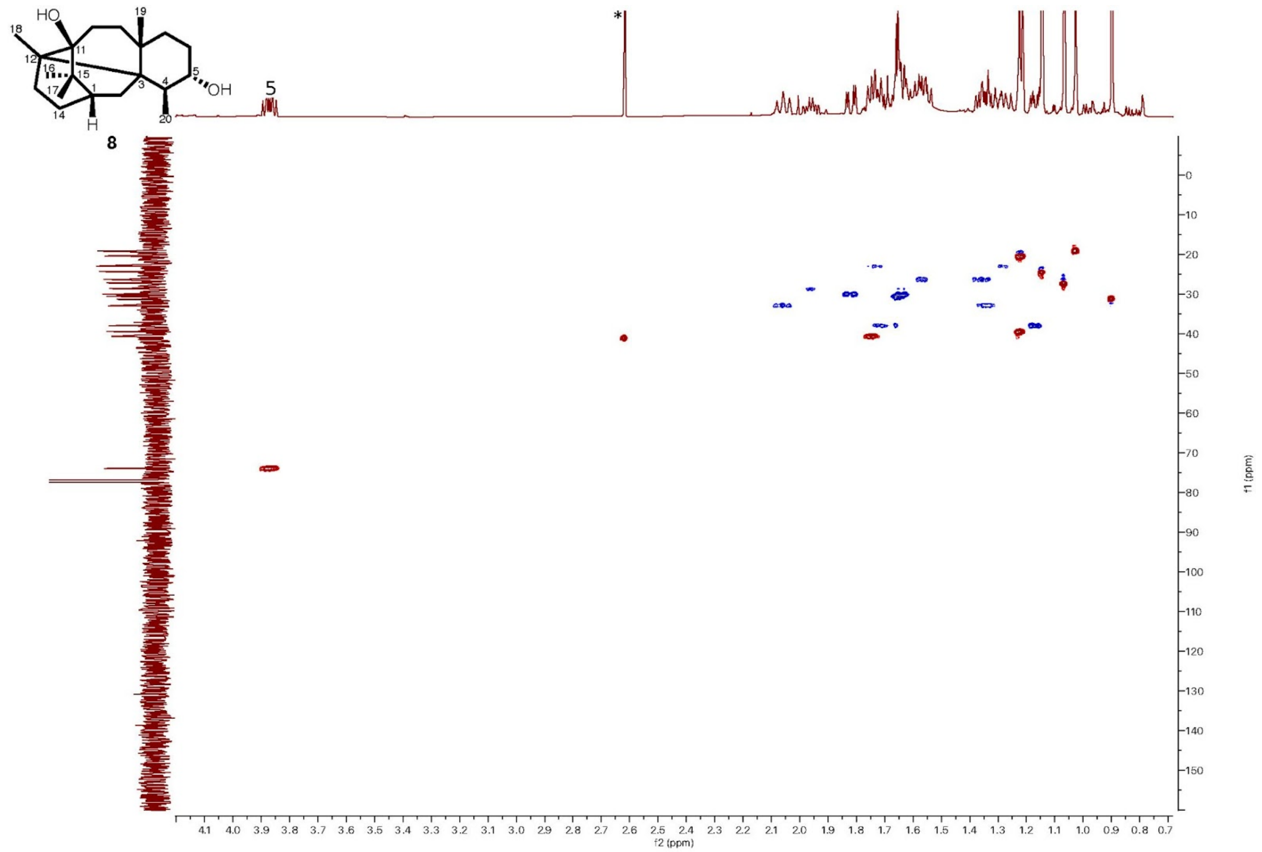


**Supplementary Figure 30. HSQC spectrum of compound 8 in $CDCl_3$ (600 Hz, n = 8).**

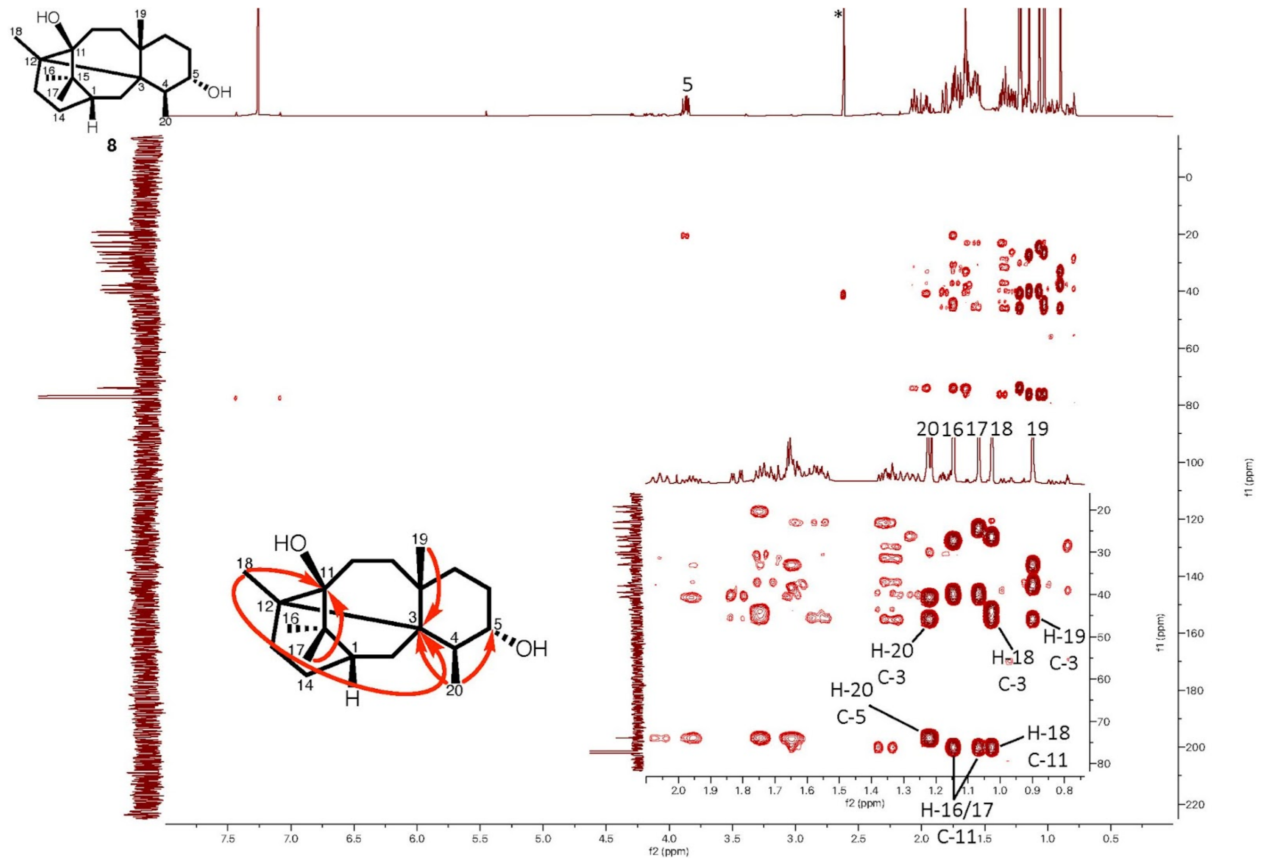


**Supplementary Figure 31. HMBC spectrum of compound 8 in $CDCl_3$ (600 Hz, n = 32).**

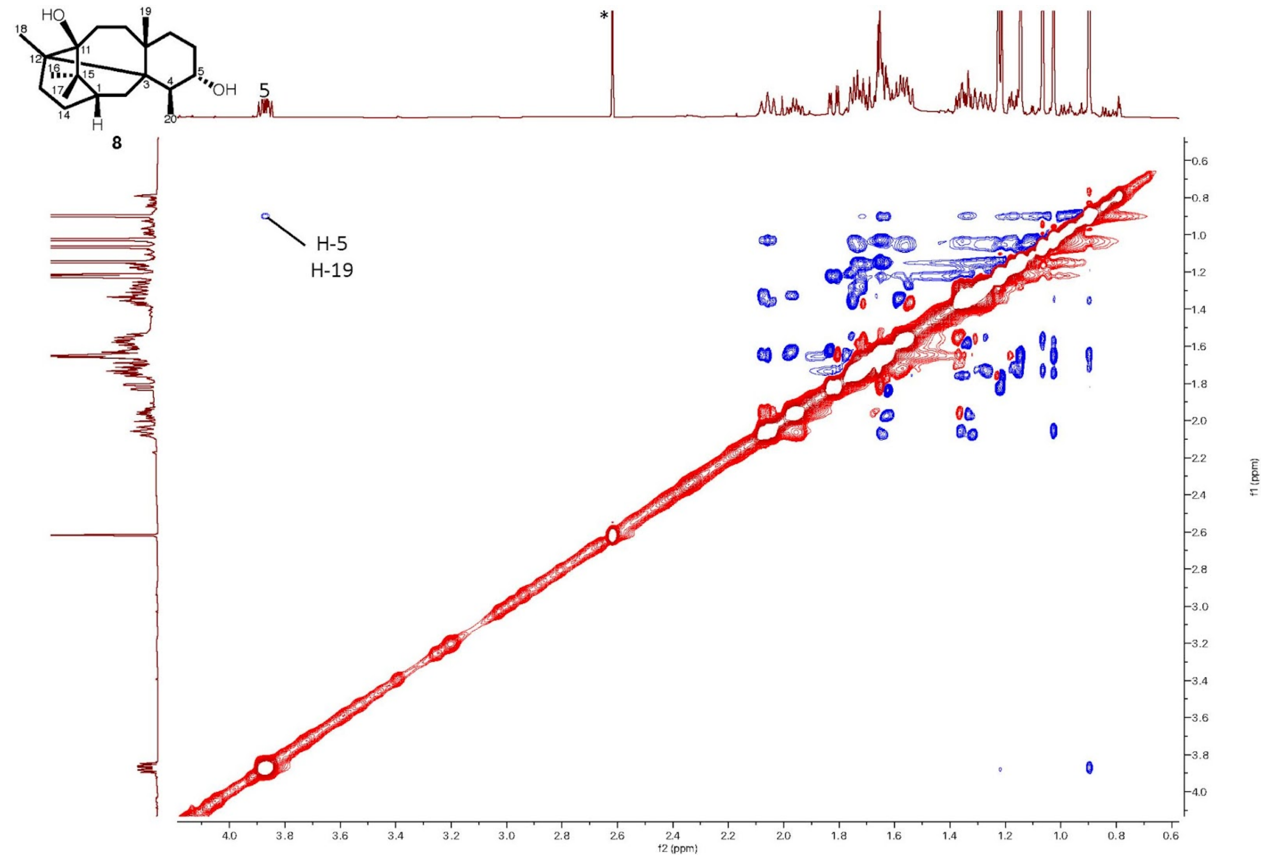


**Supplementary Figure 32. ROESY spectrum of compound 8 in $CDCl_3$ (600 Hz, n = 8).**

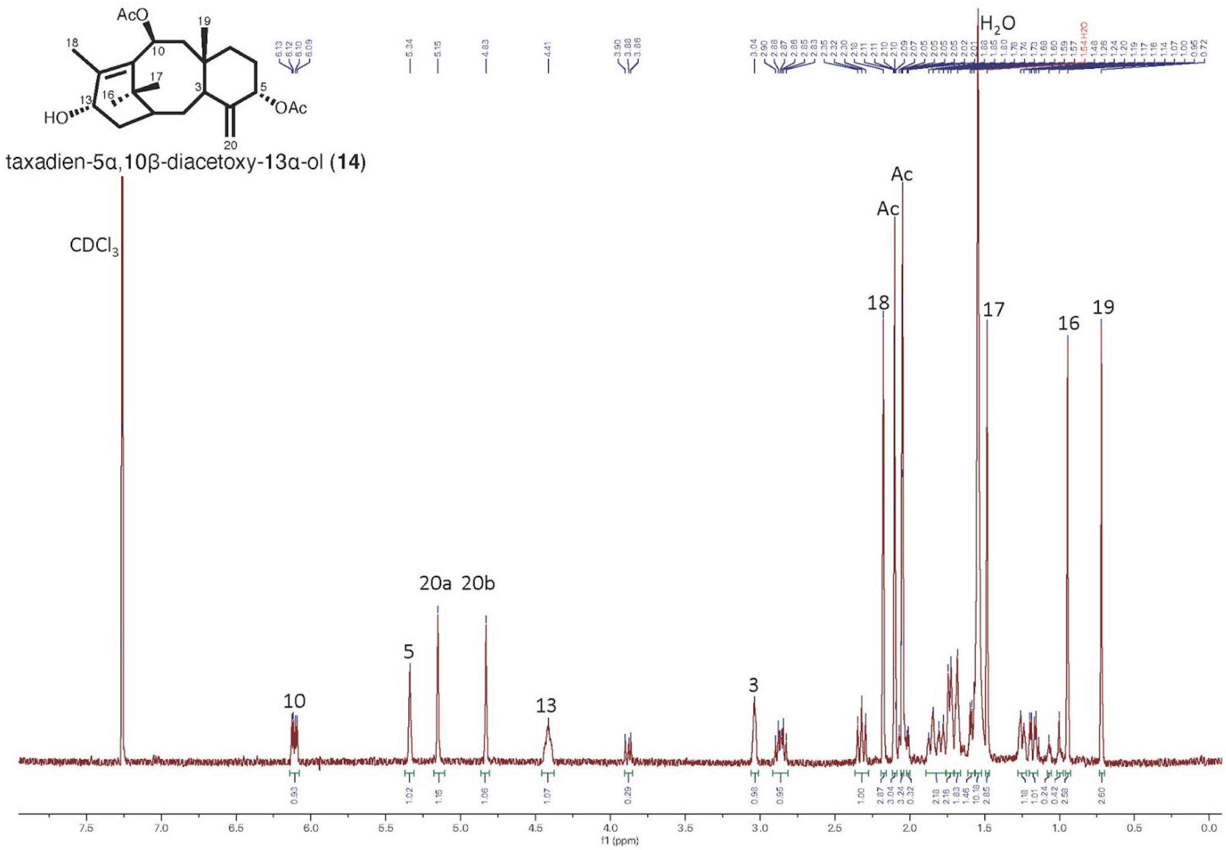


**Supplementary Figure 33.** $^{1}$**H-NMR spectrum of 5α,10β-diacetoxy-13α-ol (14) in** $CDCl_3$ **(500 Hz, n = 64).**

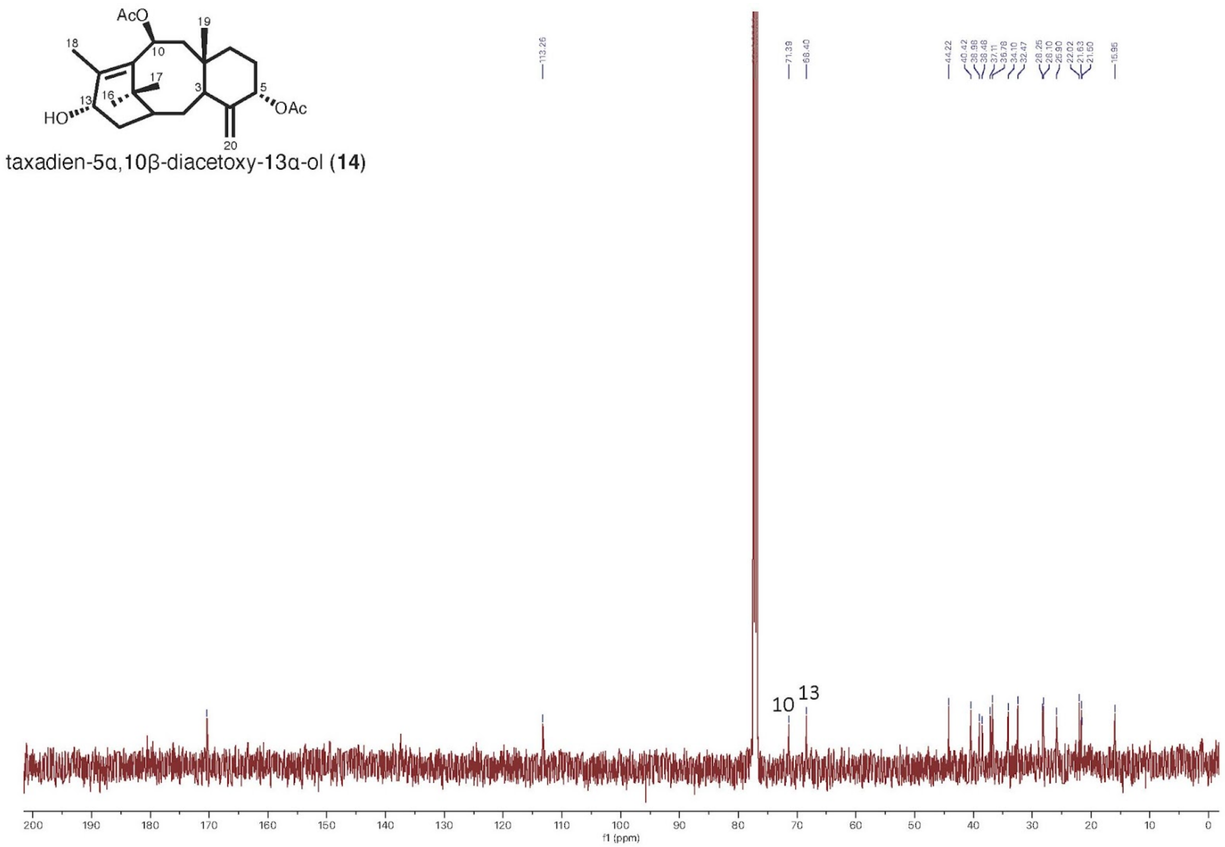


**Supplementary Figure 34. $^{13}$C-NMR spectrum of 5α,10β-diacetoxy-13α-ol (14) in $CDCl_3$ (500 Hz, n = 2048).**

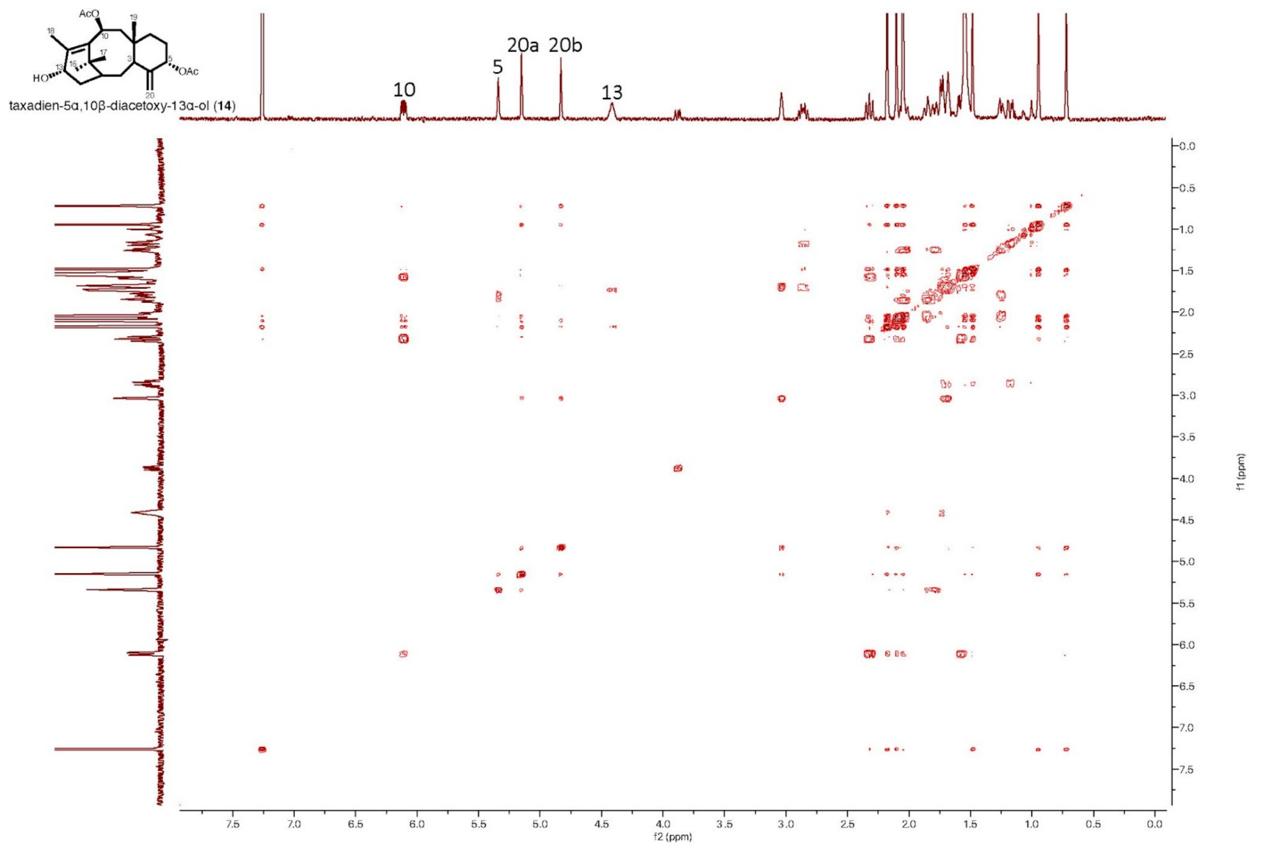


**Supplementary Figure 35. COSY spectrum of 5α,10β-diacetoxy-13α-ol (14) in $CDCl_3$ (500 Hz, n = 4).**

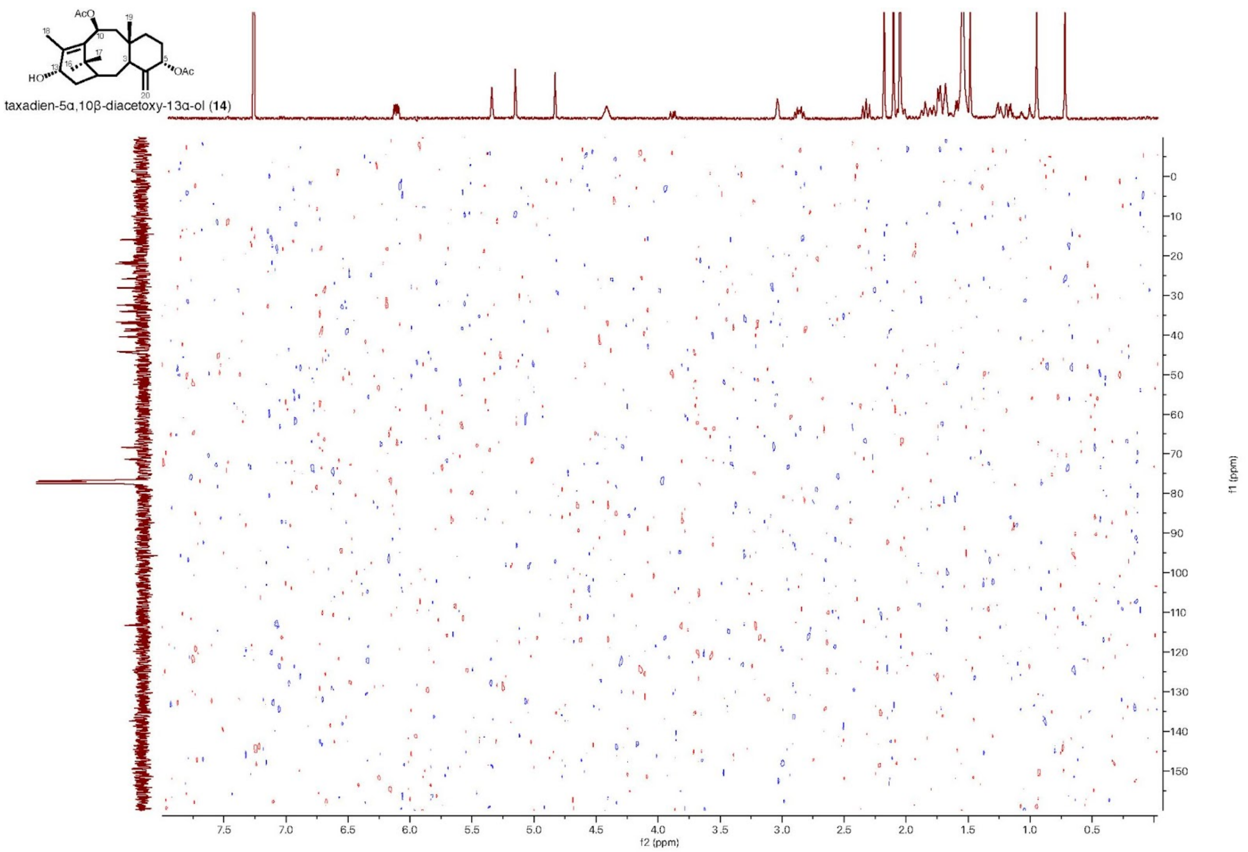


**Supplementary Figure 36. HSQC spectrum of 5α,10β-diacetoxy-13α-ol (14) in $CDCl_3$ (500 Hz, n = 8).**

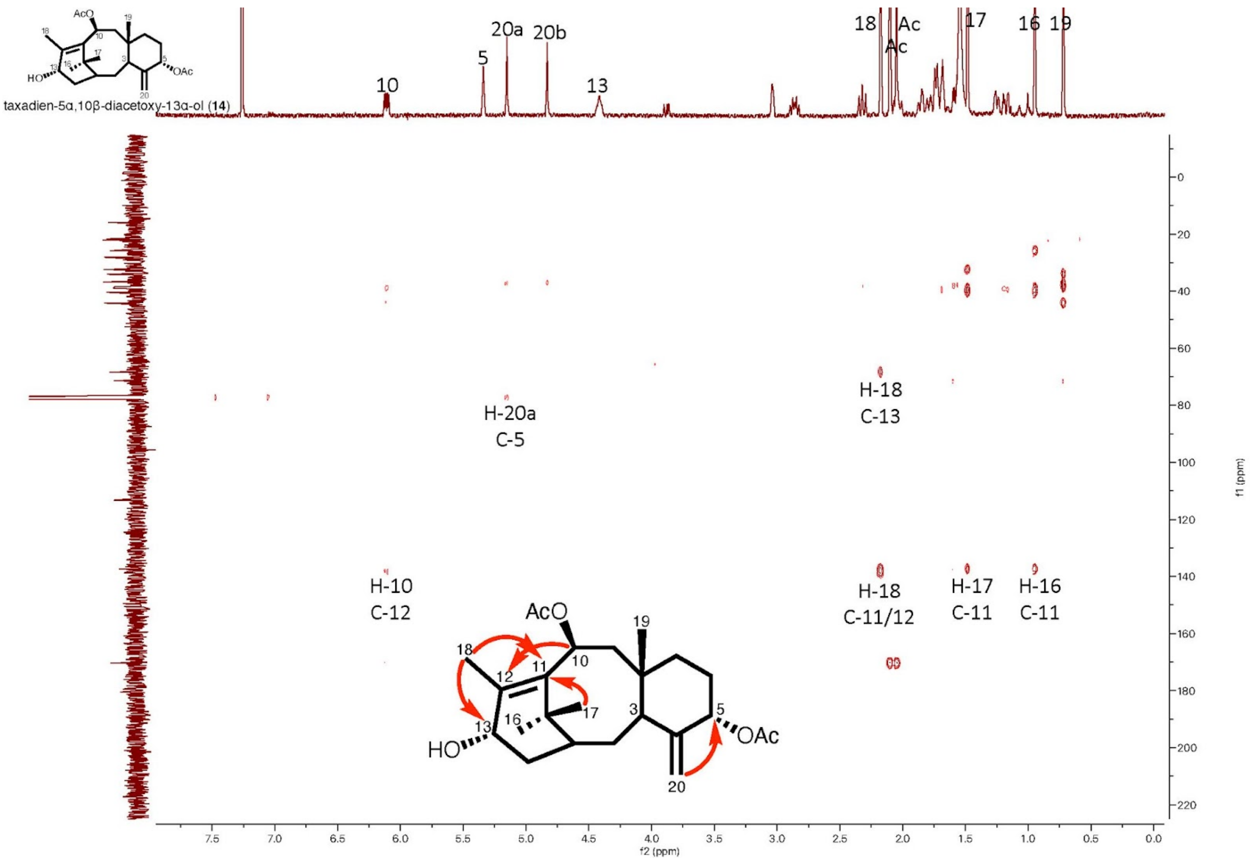


**Supplementary Figure 37. HMBC spectrum of 5α,10β-diacetoxy-13α-ol (14) in $CDCl_3$ (500 Hz, n = 32).**

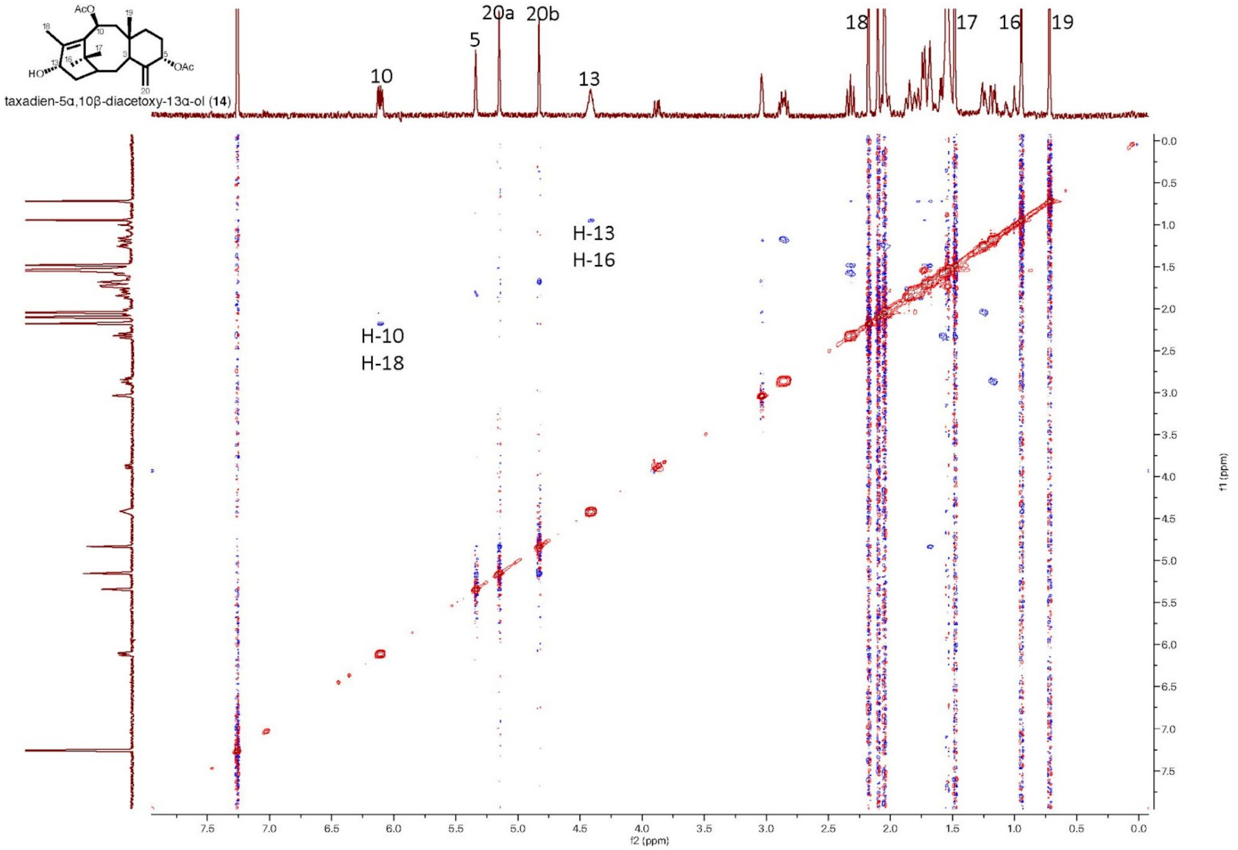


**Supplementary Figure 38. ROESY spectrum of 5α,10β-diacetoxy-13α-ol (14) in $CDCl_3$ (500 Hz, n = 8).**

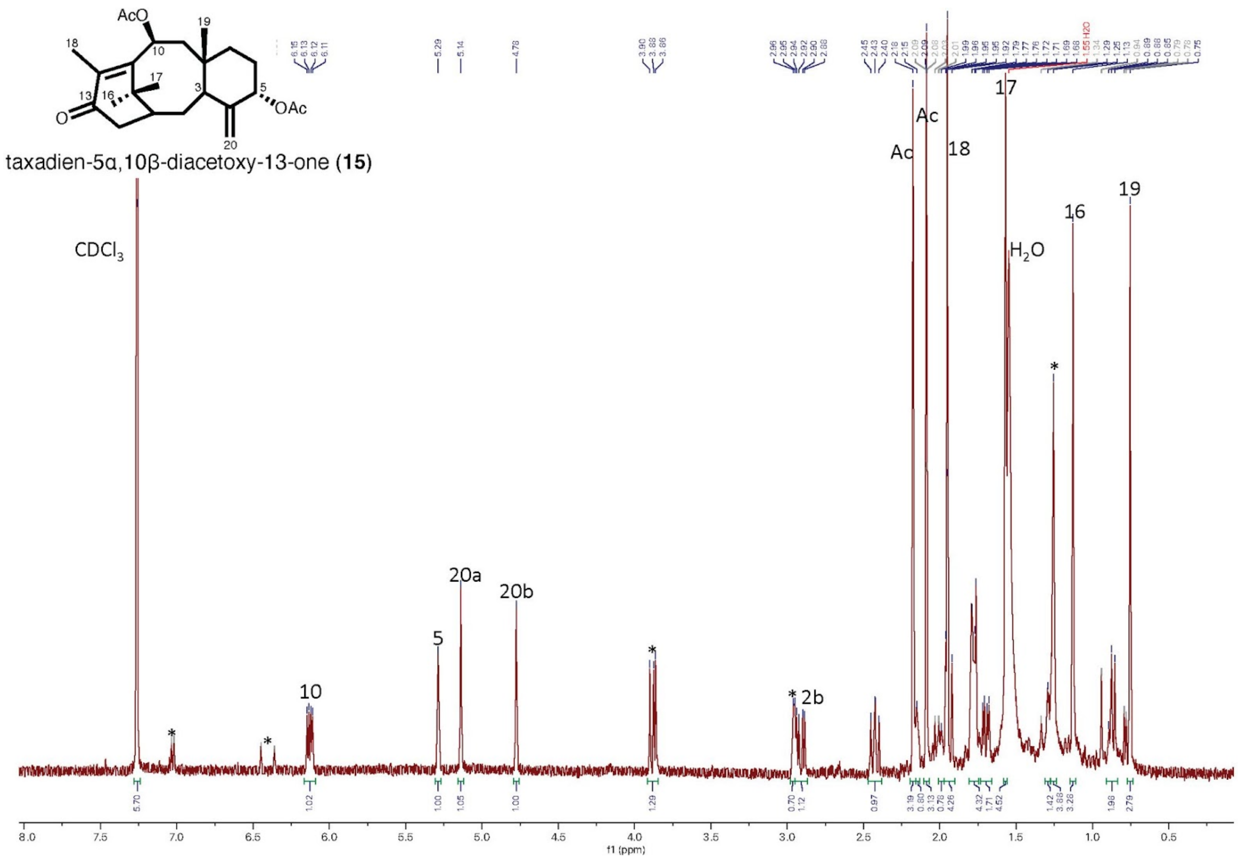


**Supplementary Figure 39. $^1$H-NMR spectrum of 5α,10β-diacetoxy-13α-one (15) in $CDCl_3$ (500 Hz, n = 64).** Asterisk indicates impurities.

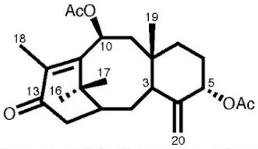


taxadien-5α,10β-diacetoxy-13-one (**15**)

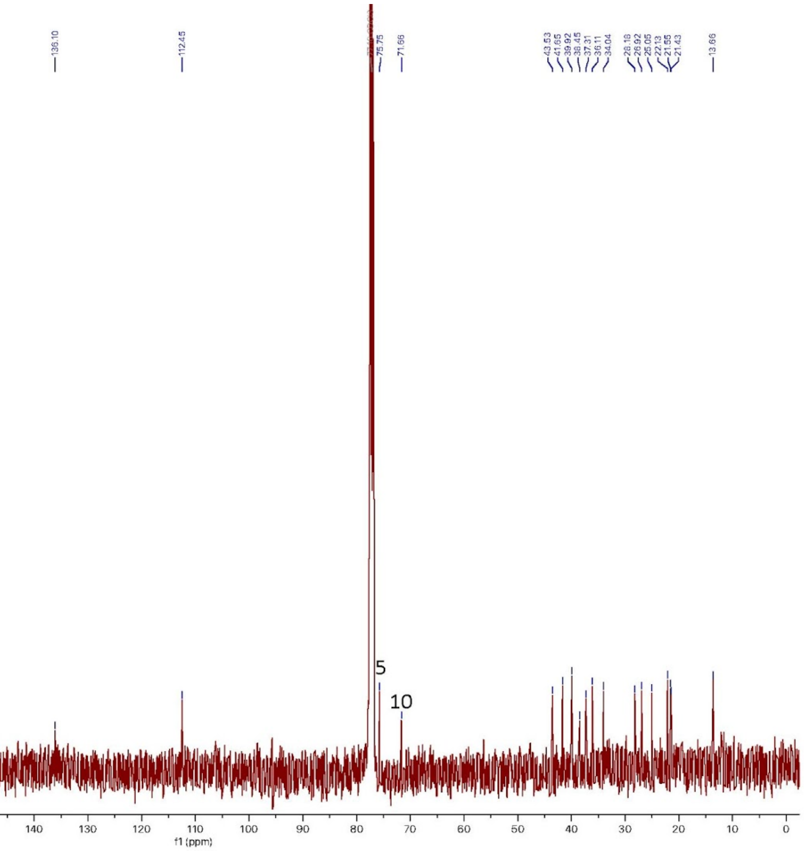


**Supplementary Figure 40. $^{13}$C-NMR spectrum of 5α,10β-diacetoxy-13α-one (15) in $CDCl_3$ (500 Hz, n = 4096).**

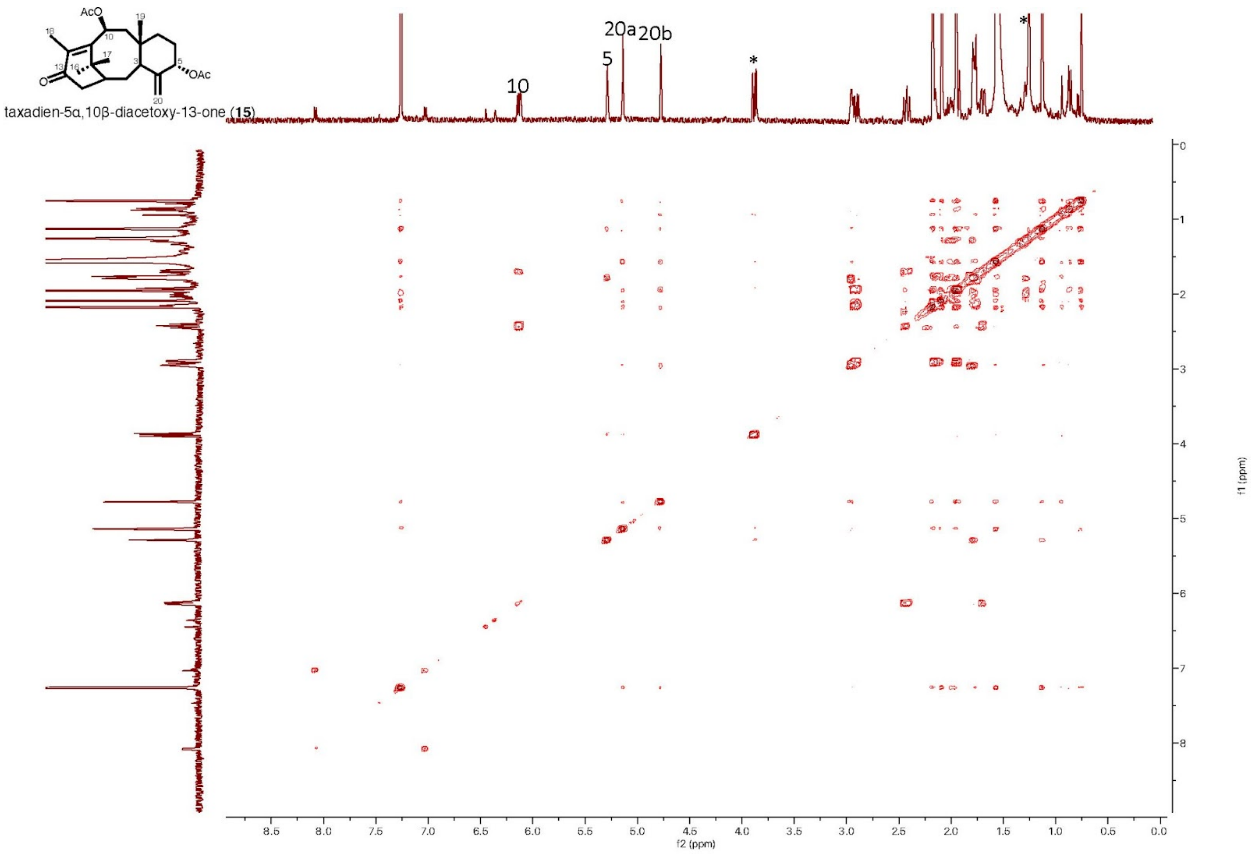


**Supplementary Figure 41. COSY spectrum of 5α,10β-diacetoxy-13α-one (15) in $CDCl_3$ (500 Hz, n = 2).**

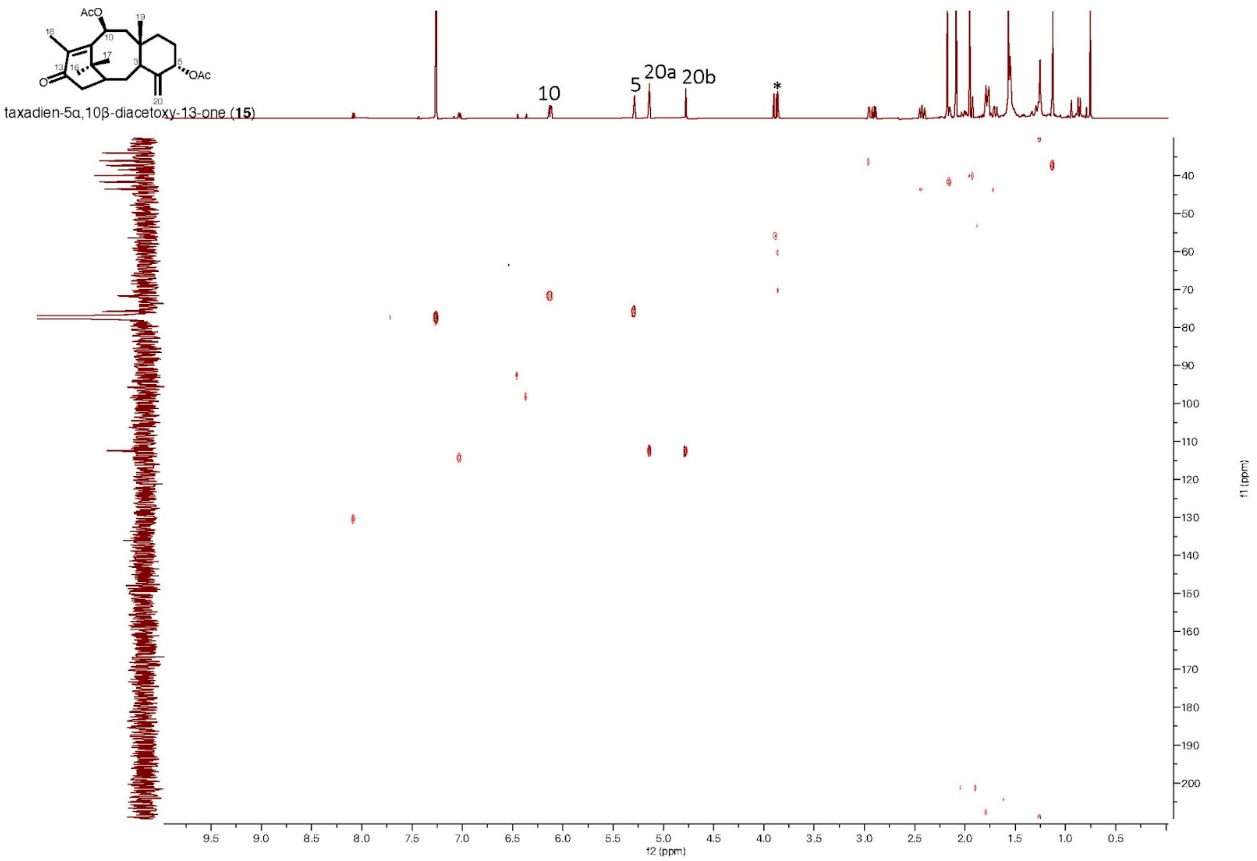


**Supplementary Figure 42. HSQC spectrum of 5α,10β-diacetoxy-13α-one (15) in $CDCl_3$ (600 Hz, n = 8).**

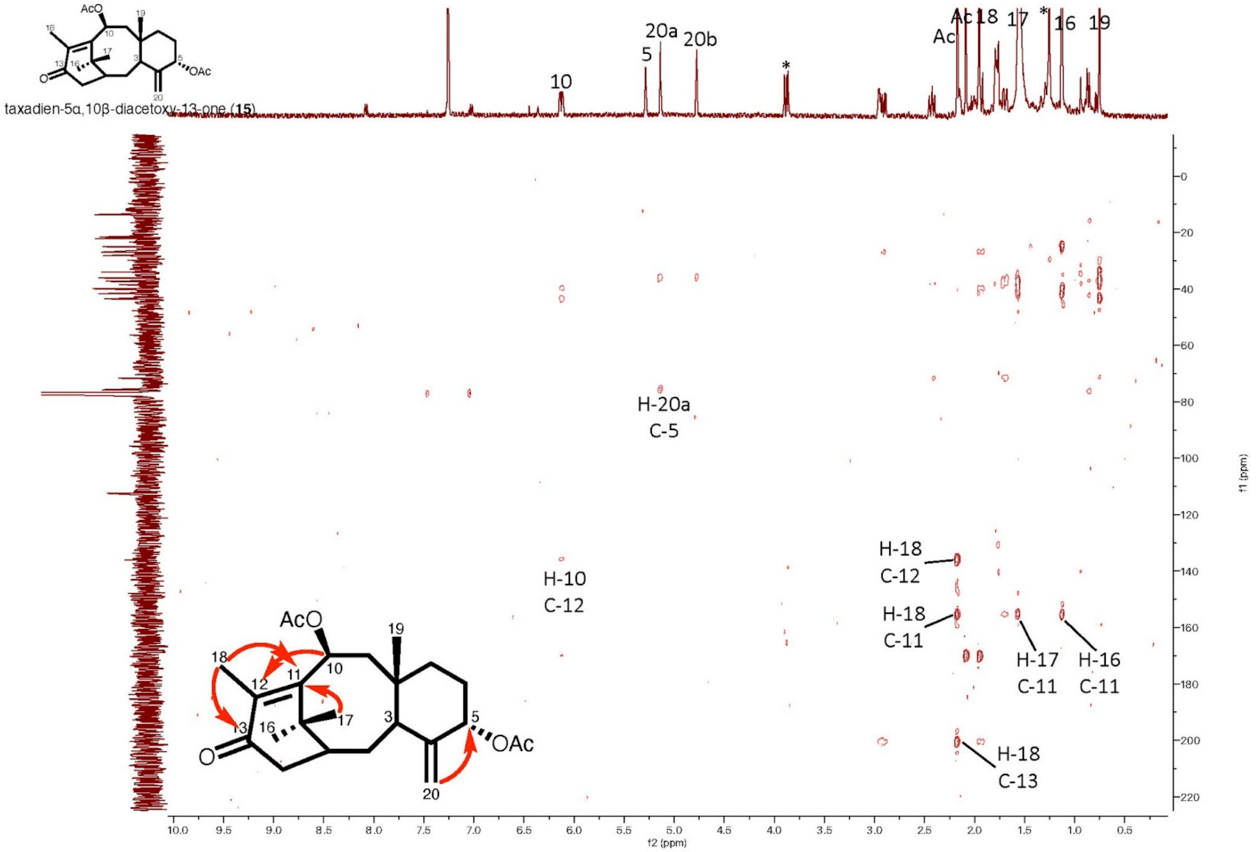


**Supplementary Figure 43. HMBC spectrum of 5α,10β-diacetoxy-13α-one (15) in $CDCl_3$ (500 Hz, n = 64).**

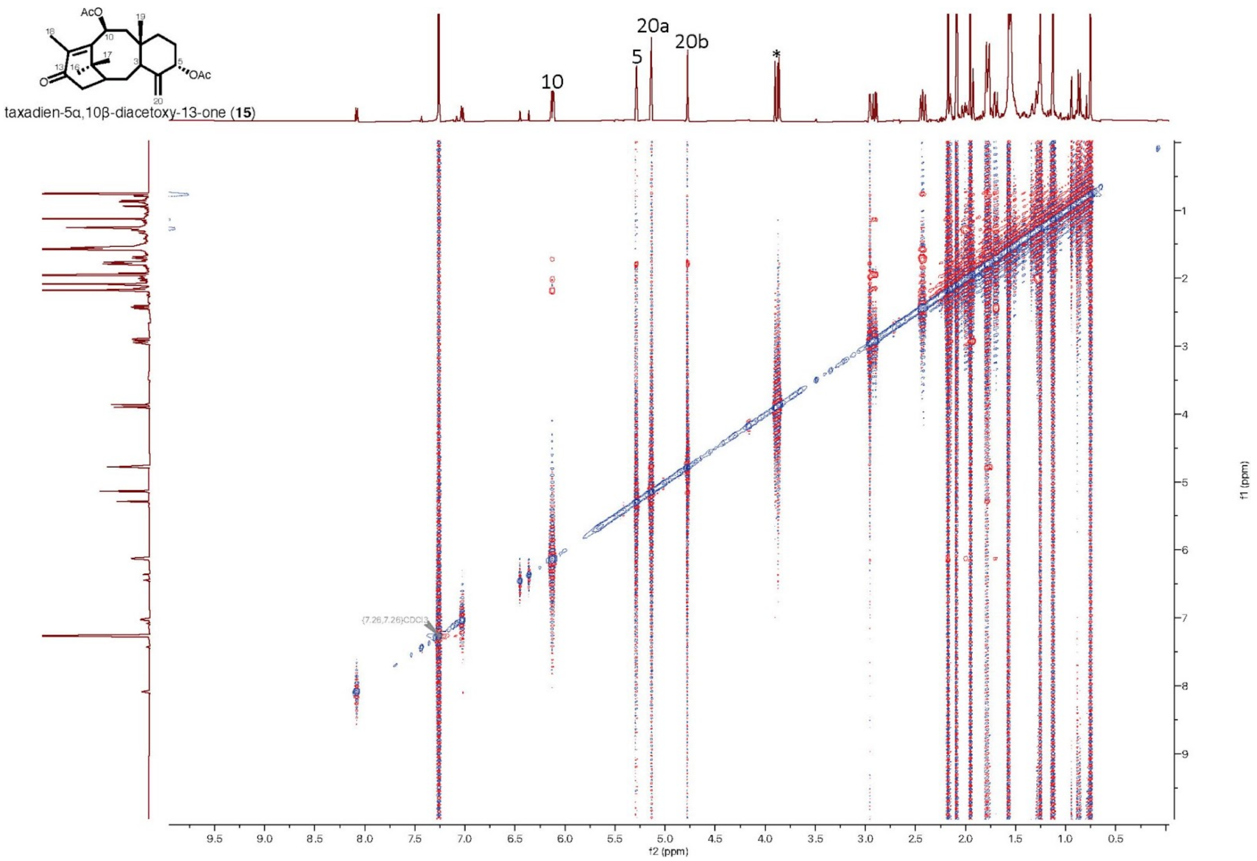


**Supplementary Figure 44. ROESY spectrum of 5α,10β-diacetoxy-13α-one (15) in $CDCl_3$ (500 Hz, n = 8).**
